# Single Cell Landscape of Sex-specific Drivers of Alzheimer’s Disease

**DOI:** 10.1101/2025.04.18.647076

**Authors:** Yiyang Wu, Kyle J. Travaglini, Mariano Gabitto, C. Dirk Keene, Amy R. Dunn, Catherine C. Kaczorowski, Philip L. De Jager, Vilas Menon, Julie A. Schneider, David A. Bennett, Logan Dumitrescu, Timothy J Hohman

**Affiliations:** Vanderbilt Memory and Alzheimer’s Center, Vanderbilt University Medical Center, Nashville, TN, USA; Vanderbilt Genetics Institute, Vanderbilt University Medical Center, Nashville, TN, USA; Allen Institute for Brain Science, Seattle, WA, USA; Department of Laboratory Medicine and Pathology, University of Washington, Seattle, WA, USA; The Jackson Laboratory, Bar Harbor, ME, USA; Department of Neurology, University of Michigan, Ann Arbor, Michigan, USA; Center for Translational & Computational Neuroimmunology, Department of Neurology, Columbia University Irving Medical Center, New York, NY, USA; Taub Institute for Research on Alzheimer’s Disease and the Aging Brain, Columbia University Irving Medical Center, New York, NY, USA; Rush Alzheimer’s Disease Center, Rush University Medical Center, Chicago, IL, USA

**Keywords:** Alzheimer’s disease, Dementia, Sex difference, Sex specific, Single cell, RNA sequencing, Transcriptome, Signaling pathway

## Abstract

**Background:** Sex differences in Alzheimer’s disease (AD) have been documented for decades, and many sex-specific molecular contributors to AD have been discovered through bulk omics analysis of brain tissues. RNA sequencing (RNAseq) at single cell resolution provides an opportunity to characterize transcript associations with AD in a cell type-specific matter. Here, we investigated sex-specific gene expression associations with neuropathology and cognitive manifestation of AD (endophenotypes) leveraging a large single-nucleus transcriptomic dataset consisting of 1.64 million nuclei from dorsolateral prefrontal cortex (DLPFC) tissue of 424 unique donors from the Religious Orders Study and Memory and Aging Project (ROS/MAP; AD Knowledge Portal syn2580853).

**Methods:** ROS/MAP single-nucleus RNAseq data (snRNA-seq) were processed through a rigorous pipeline. In total, eight major cell types from DLPFC were identified. We first performed sex-stratified and sex-interaction association analyses by fitting negative binomial mixed models in relation to β-amyloid load (Aβ), paired helical filament tau tangle density (tau), global cognitive performance at last visit, and longitudinal cognitive trajectory. We then conducted gene-set enrichment analysis to identify functional signaling pathways enriched for sex-specific associations. Lastly, we compared differential gene expression patterns and intercellular communication profiles between sexes and diagnostic groups among major cell types. For replication, sex-specific associations were examined using snRNA-seq derived from DLPFC tissue-derived of an independent set of 84 donors from The Seattle Alzheimer’s Disease Brain Cell Atlas (SEA-AD) study.

**Results:** 68% of the ROS/MAP participants were female, and 52% were diagnosed with AD dementia. We first identified several disease-dependent or sex-dependent cell subpopulations. Then we identified 2,660 sex-specific associations involving 2,110 genes with Aβ (51%), tau (21%), and cognitive performance (29%). 60% female-specific associations were for Aβ, and 49% male-specific associations were with tau. The vast majority (93%) of female protective associations were from neurons, and most (76%) of female risk associations were from glial cells. Nine of the female-specific associations involving eight unique genes were replicated in the SEA-AD cohort, including *ADGRV1* and *OR3A3* with Aβ; *IFI27L1*, *LYRM1*, *STAP2*, and *TSTD2* with tau; *PDYN* with global cognition; and *TMEM50B* with longitudinal cognitive decline. All replicated associations except *TMEM50B* were observed in neurons. Furthermore, the preponderance of protective female-specific associations in neurons was also recapitulated in the SEA-AD cohort. Sex-specific associations were enriched for genes in the immune, inflammation, and damage-related stress response pathways, and microglia presented the most sex-specific enriched pathways. Finally, we identified six ITGB1-mediated microglia-specific incoming signals that may play a role in female-specific risk for Aβ accumulation.

**Conclusion:** Our study highlights the transcriptome-wide, single-cell landscape of sex-specific molecular associations with AD neuropathology and cognitive decline. We delineate the full scope of sex-specific transcript associations, differential expression, signaling pathway, and cell-cell communication network changes in each major DLPFC cell type, while identifying and replicating several female-specific gene associations in neurons to help direct future mechanistic studies.

## Background

Alzheimer’s disease (AD) has a higher prevalence in women, with women making up two-thirds of prevalent AD cases^1,2^. While differences in life-span contribute to the gender disparity in AD, several studies have shown that even at the same age, women are more likely to develop AD than men^2,3^. Women with AD also tend to have more global AD pathology^4^, faster cognitive decline^5,6^, faster brain atrophy^7^, and experience enhanced risk when carrying the *APOE*-ε4 allele^8–10^. In contrast, men with AD have a higher mortality rate and a greater comorbidity burden^11,12^. Thus, there is a pressing need to better characterize the molecular contributors that underlie such sex differences in the neuropathology and clinical manifestation of AD.

Our team has been particularly interested in identifying genetic factors that act in a sex-specific manner to modify risk and resilience in AD. We and others have identified novel female-specific and male-specific drivers of AD neuropathology^13,14^, cognitive decline^15^, and resilience to cognitive decline leveraging advanced genomic approaches^16^. Additionally, we have provided a deep characterization of sex-specific associations between *APOE* and AD neuropathology^17^ and cognitive decline^18^. Together, our work has suggested that the sex-specific genetic drivers of AD emerge largely downstream of Aβ with a notable sex-specific genetic architecture that contributes to tau deposition, neurodegeneration, and cognitive decline^19^.

In addition to the genomic evidence, multiple studies have characterized sex differences in AD using bulk multiomics measured in brain and blood tissue of humans and mice^20–23^, consistently highlighting that sex-specific genes are enriched for immune response. However, most of these studies either focused on roles of sex chromosomes and sex hormones or were limited to differentially expressed genes (DEGs) between sexes. The advancement of single cell/nucleus RNA sequencing in recent years has facilitated a more complete dissection of the molecular mechanisms that contribute to sex differences in AD at single-cell resolution. One study characterized sex differences in AD neuropathology and cognitive decline at the single-cell level leveraging data generated from 48 brains in the Religious Orders Study and Memory and Aging Project (ROS/MAP), highlighting multiple female-specific protective associations in neurons and male-specific risk associations in oligodendrocytes^24^. Our study builds on this important work by leveraging a much larger single-nucleus transcriptomic dataset (AD Knowledge Portal syn2580853) derived from dorsolateral prefrontal cortex (DLPFC) tissue of 424 donors from ROS/MAP^25^ and performing an independent replication of sex-specific associations utilizing another single-nucleus RNA sequencing (snRNA-seq) dataset from 84 donors in The Seattle Alzheimer’s Disease Brain Cell Atlas (SEA-AD) study^26^. The goal of this manuscript is to comprehensively characterize the transcriptome-wide single cell landscape of sex-specific molecular associations with AD neuropathology and cognitive decline. We hypothesized that sex-specific effects would emerge in a cell-type specific manner, and based on studies mentioned above, that we would observe female-specific protective associations in neurons, male-specific risk associations in oligodendrocytes, as well as sex-specific associations enriched in immune cells. Finally, we move beyond single-transcript associations to provide detailed cell-cell communication alterations in the AD brain that act in a sex-specific manner, along with biological pathways that show sex-specific enrichment in the AD brain.

## METHODS

### Participants

The participants included in this study were from two longitudinal clinical-pathological cohort studies including the Religious Orders Study and the Rush Memory and Aging Project (ROS/MAP). Each study was approved by the Institutional Review Board (IRB) of Rush University Medical Center. At enrollment, all participants were free of known dementia and agreed to annual clinical evaluation and brain donation^27–29^. All participants signed an informed repository consent and an Anatomic Gift Act. Secondary analyses of this extant data were approved by the Vanderbilt University Medical Center IRB.

Independent replication was sought leveraging snRNA-seq data from 84 donors from The Seattle Alzheimer’s Disease Brain Cell Atlas (SEA-AD) study^30^. Postmortem brain tissue and donor metadata of SEA-AD study were obtained via the University of Washington (UW) BioRepository and Integrated Neuropathology (BRaIN) laboratory from participants in the Kaiser Permanente Washington Health Research Institute Adult Changes in Thought (ACT) Study and the University of Washington Alzheimer’s Disease Research Center (ADRC). Informed consent for research brain donation was obtained according to protocols approved by the UW and Kaiser Permanente Washington Health Research Institute IRBs.

There are various fundamental differences underlying these two cohorts, including cohort characteristics, sequencing data generation, quality control process, and cell type taxonomies that we discuss in detail in the limitations section of the discussion.

### Single-nucleus RNA Sequencing

Single-nucleus transcriptomes of DLPFC brain specimen derived from 465 ROS/MAP participants (AD Knowledge Portal Accession Number syn31512863) were collected by the Rush ADRC and processed at Columbia University Medical Center, including sample processing, library preparation and sequencing of single nuclei, processing of single-nucleus RNAseq reads, quality control, cell type classification, and cell sub-classers analysis. Detail methods can be found in the previous report^31^. Post-quality control (post-QC) datasets contained 424 unique participants with a median of 3,824 sequenced nuclei. Eight major cell types were clustered and used for analysis, including astrocytes, vascular niches (referred to as “endothelial cells” in this article), microglia, oligodendrocytes, oligodendrocyte precursor cells (OPCs), inhibitory neurons, and *CUX2+* and *CUX2-* excitatory neurons (Fig. 1b). Excitatory neurons were split into two subgroups to increase computational efficiency (*CUX2+* consisted of upper cortical layer 2-4 pyramidal neurons, and *CUX2-* consisted of all others deeper cortical layer pyramidal neurons). Genes with expression in at least 10% of all cells were used for downstream analyses. Cells were removed from downstream analyses if they counted more than 20,000 or less than 200 total RNA Unique molecular identifiers (UMIs) or had more than 5% mitochondrially-mapped reads. RNA molecular count data was normalized and scaled by the “sctransform” R package^32^ (v2) using “percent.mt” values for “vars.to.regress” parameter.

**Fig. 1:**
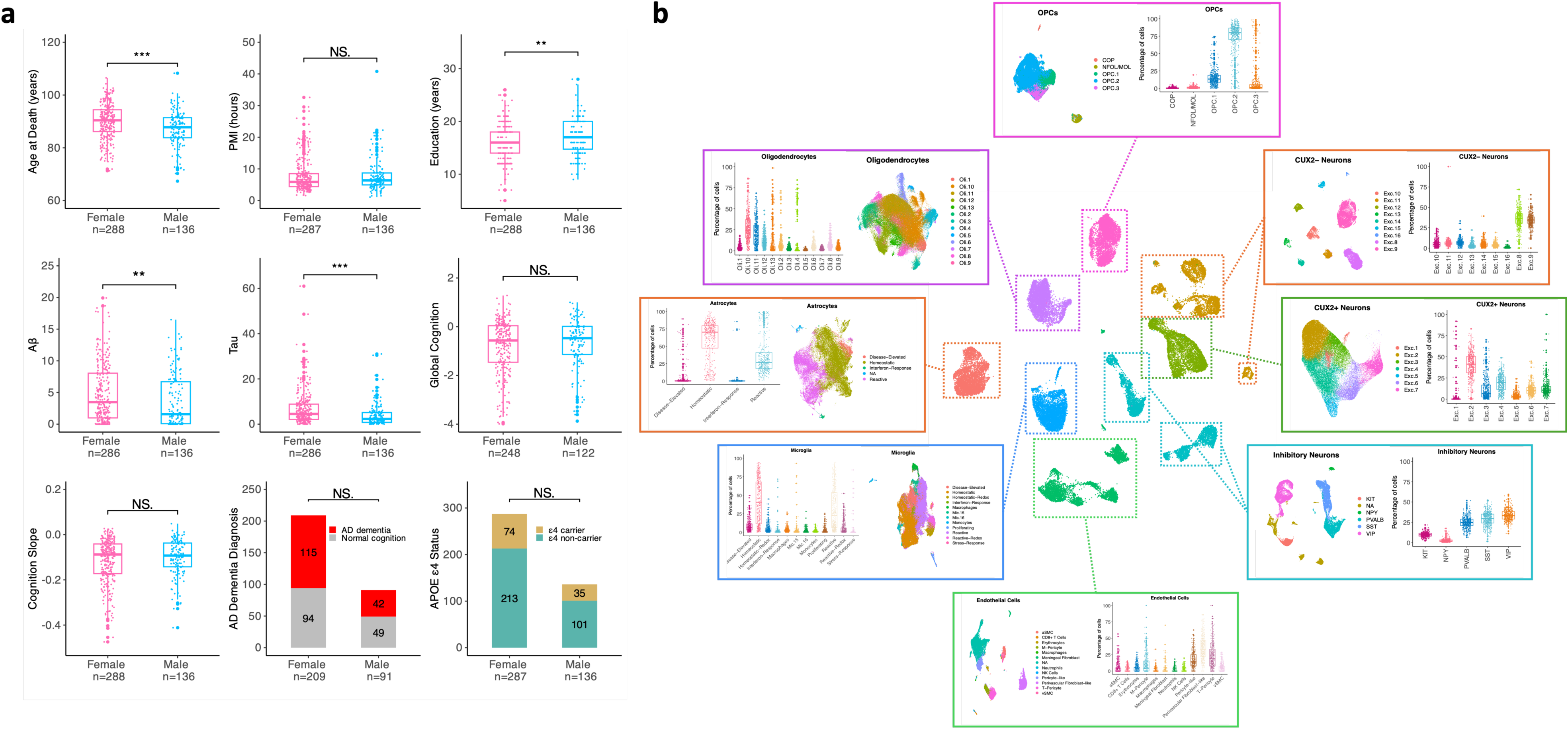
ROS/MAP snRNA-seq cohort characteristics and cell type annotations. a, ROS/MAP snRNA-seq cohort characteristics summary, showing age at death, PMI, education length, Aβ, tau, global cognition at last visit, cognition slope, AD dementia diagnosis, and APOE ε4 carrier status. ***p<0.001, **p<0.01, *P<0.05, Abbreviations: NS, not significant. PMI, post-mortem interval. For APOE ε4 carrier status and diagnosis, statistical testing was done for the number of ε4 carriers or AD dementia patients between sexes. b, Joint uniform manifold approximation and projection (UMAP), colored by major cell type. Composition of each major cell type was represented by subtype UMAP and box plots.

The snRNA-seq dataset of SEA-AD (syn52146347) consisted of 1.2 million nuclei derived from 84 donors’ postmortem DLPFC (Brodmann area 9) tissues. Cells were mapped to the NIH’s Brain Research through Advancing Innovative Neurotechnologies (BRAIN) Initiative - Cell Census Network (BICCN)-based cellular taxonomy^33^, which resulted in 24 subclasses including LAMP5 LHX6, LAMP5, PAX6, SNCG, VIP, SST CHODL, SST, PVALB, CHANDELIER, L2/3 IT, L6 IT, L4 IT, L5 IT, L5 ET, L6 CT, L6B, L6 IT Car3, L5/6 NP, astrocytes, OPC, oligodendrocytes, endothelial cells, VLMC, and Microglia-PVM.

### Cell Composition Analysis

Cell composition analyses of the ROS/MAP snRNA-seq dataset include both cell fraction comparison and cell abundance comparison between sexes and AD diagnostic groups for the eight major cell types as well as for the 67 subgroups (unidentified “NA” (unassigned) subgroups were removed). Cell fraction (percentage) was calculated with “NA” subgroups included and compared by two-way ANNOVA with sex-by-AD dementia diagnosis interaction term. Cell abundance analyses were carried out using “crumblr” (v0.99.11) R package (https://DiseaseNeurogenomics.github.io/crumblr). For cell abundance analyses, subpopulation cell counts were centered log ratio (CLR) transformed using “crumblr” function, which also returned observation-level inverse-variance weights for use in weighted linear models to test compositional change of each cell type separately following the formula: ∼ age at death + postmortem interval (PMI) + sex + AD dementia diagnosis + sex*AD dementia diagnosis implemented by the “dream” function in the “crumblr” package. 299 participants were included in this analysis after removing those without PMI or AD dementia diagnosis data. P-values were adjusted using a Bonferroni correction across all 8 cell types or 67 subgroups tested.

### AD Endophenotype Measurements

For the ROS/MAP cohorts, global β-amyloid load (Aβ, “amyloid” variable from Research Resource Sharing Hub (RADC)) and paired helical filament tau tangle density (tau, “tangles” variable from RADC) were measured at autopsy by immunohistochemistry staining, quantified as the average percent area occupied by Aβ (using antibodies specific to Aβ42) or tau (using antibodies specific to AT8 epitope of abnormally phosphorylated tau) across eight brain regions at autopsy: hippocampus, angular gyrus, and entorhinal, midfrontal, inferior temporal, calcarine, anterior cingulate, and superior frontal cortices. Then, values were transformed to approximate a normal distribution^34^.

The global cognitive score was calculated as a composite measure of global cognitive function by converting raw scores from 17 cognitive tests to Z scores and calculating their average^35^. For cross-sectional analyses of cognitive function, the global cognitive score at last visit before death was used. Due to the mixed effects model required for snRNA analysis, longitudinal cognitive trajectory was evaluated by quantifying a cognitive slope derived leveraging a linear mixed effects model with global cognition as the outcome, the intercept and years from last cognitive visit entered as both fixed effects and random effects. The random slope is the estimated person-specific rate of change in the global cognition variable over time. This person-specific estimate was added to the fixed effects slope to calculate each individual’s annual rate of change.

For the SEA-AD DLPFC cohort, we used gold standard Consortium to Establish a Registry for Alzheimer’s Disease (CERAD) neuritic plaque staging (none, sparse, moderate, frequent) and Braak neurofibrillary tangle staging (stages 0-VI) and as semi-quantitative outcomes to represent Aβ and tau pathology, respectively, due to lack of quantitative pathology scores during the preparation of this manuscript. Slopes of memory decline over time based on the original Cognitive Abilities Screening Instrument (CASI) scores (to derive stable and generalizable estimates) were calculated using mixed-effects models with an unstructured covariance matrix, where time was parameterized as years before death. The CASI score at the last visit was used to represent global cognition at last visit.

### Sex-specific Association Analyses with AD Endophenotypes

Gene expression associations using the ROS/MAP dataset with four AD endophenotypes (global cognition longitudinally and at last visit before death, Aβ, and tau at death) and AD dementia diagnosis were conducted for each gene in each of the eight major cell type using a negative binomial mixed model implemented in the R package “Nebula” (v1.4.2).^36^ The number of genes analyzed in each association model ranged from 14,537 to 16,672. Only participants with a PMI less than 24 hours were included, which yielded 417 participants.

To estimate associations with Aβ, tau, global cognition at last visit before death, and AD dementia diagnosis, the following formula was used; “gene count matrices ∼ AD endophenotype + sex + age at death + PMI”. For analyses of longitudinal cognitive performance, interval years between last visit and death was added as covariate together with others mentioned above. Four participants who had more than five years between their last visit and death were removed from this analysis. The gene count matrices used in all association models were taken from the “count” slot of the SCT assay (transformed RNA count assay generated by “sctransform” package mentioned above) from the Seurat object associated with each major cell type. All models were run in a combined-sex and sex-stratified manner. To estimate the sex-interaction effect with each AD endophenotype, an interaction term of “AD endophenotype*sex” was added to each model described above with the same covariates. Lowly-expressed genes were removed from the analysis in Nebula be default setting (counts per cell < 0.5% and number of cells with a positive count < 5) to obtain accurate estimates of the overdispersions. Finally, association models from Nebula with convergence scores less than or equal to -20 were removed from downstream analysis.

Sex-specific association analysis of the SEA-AD dataset was performed using the same model setup and inclusion criteria above. We carried out the association analyses in each of the pre-defined 25 cell subclasses (L2L3 subclass was split into two separate files for analysis due to computational limitation) and then reclassified them into seven major cell types to match ROS/MAP cell taxonomy for comparison. Only gene-phenotype association pairs that showed consistent direction of effect across all subclasses under the same major cell type were reserved for comparison.

Sex-specific associations of ROS/MAP were defined as those which had nominal p values < 0.05 from sex-interaction models and had false discovery rate (FDR)-adjusted p-values < 0.05 from sex-stratified models and showed the same direction of effect on the same AD endophenotype in the same cell type tested.

### Differential Gene Expression Between Sexes and AD Dementia Diagnosis

Differential gene expression (DGE) analyses in each of the eight major cell types of the ROS/MAP dataset among participants with AD dementia or normal cognition were carried out using “Nebula” R package as described above with the same covariates. DGE analysis was set up to detect gene expression changes not only within the same sex cross diagnostic groups, but also between sexes within the same diagnostic group for completeness.

We used the final consensus cognitive diagnosis (“cogdx” variable from RADC) to categorize participants into AD dementia group or normal cognition group. In total, 299 participants had cogdx scores, among which 142 with “cogdx” = 1 were assigned to the normal cognition group, and 157 participants with “cogdx” = 4 or “cogdx” = 5 were assigned to the AD dementia group.

### Gene Set Enrichment

The gene set enrichment analysis of gene lists generated from sex-stratified association models was conducted using “fgsea” function from “fgsea” R package^37^. We used 20,354 unique genes that were detected in all cell types in post-QC ROS/MAP snRNA-seq cohorts as the background set for testing to avoid bias. We ranked all genes from each sex, cell type, and AD phenotype combination by statistical significance using ranking values calculated by -log10({p-value})*sign({log value of Fold Change, logFC}). Under this rank metric, up-regulated (logFC > 0) genes with relatively small p-values appear at the top of the list and down-regulated (logFC > 0) genes with small p-values at the bottom. We removed genes that have a ranking of infinity due to having very small p-values. Then we tested each of the 64 pre-ranked gene lists (four AD phenotypes*eight cell types*two sexes) against 50 signature gene sets in HALLMARK from MSigDB^38^ (v2023.1) with 1,000,000 permutations. P-values of enrichment tests were adjusted using Bonferroni correction cross 3,200 tests for HALLMARK gene sets.

### Intercellular Communication Profiling

Intercellular communication patterns among seven major cell types (merging both excitatory neuron subgroups) was analyzed using the “CellChat” (v2.0) R package^39^ with default setting without projecting gene expression data onto human protein-protein interaction network. ANOVA (for more than two groups) and Wilcoxon rank-sum (between two groups) tests were used for statistical comparison. Turkey HSD tests post ANOVA were run to detect difference between any two groups.

### Gene ratio calculation

To examine whether sex-specific genes located on sex chromosomes were enriched comparing to those from autosomes, we calculated gene ratio by dividing the total annotated gene number on each chromosome based on Human Genome Assembly GRCh38.p14^40^.

### Statistical Analysis

Statistical analyses were conducted in RStudio (R version 4.3.1). Significance was set *a priori* to α=0.05. P-values from all sex-stratified models were corrected for multiple comparisons using the FDR procedure across all 1,003,028 tests combining all four endophenotypes for ROS/MAP. Overlapping sex-specific associations between SEA-AD and ROS/MAP from sex-stratified models of SEA-AD were corrected for 2,660 tests using R function “p.adjust” with “BH” as the method.

## RESULTS

For the purpose of summarizing, we defined an association is protective or has a protective effect if in which a higher gene expression is associated with a better AD-related endophenotype (less neuropathology, or better cognition performance), and defined an association is risk or has a risk effect if in which a higher gene expression is associated with a worse AD-related endophenotype.

### ROS/MAP cohorts characteristics

The post-QC ROS/MAP snRNA-seq dataset was derived from 424 Non-Hispanic White donors, with a mean age at death of 89 years, and an average of 16 years of education. 68% were females, 52% participants met clinical AD dementia criteria, and 26% carried at least one *APOE*-ε4 allele. Females and males differed statistically in their age at death, education, Aβ, and tau, with females presenting at an older age at death, with less education, and higher Aβ and tau than males (Fig. 1a, Supplementary Table 1).

### Identifying vulnerable cell subpopulations in AD dementia

We first compared whether the composition of the eight major cell types in the ROS/MAP snRNA-seq dataset (Fig. 2a) was influenced by sex, AD dementia, or their interaction to provide a foundation for our explorations into the sex-specific transcript associations with AD endophenotypes. We observed no major cell type constitution differences in either cell fraction or cell abundance by sex or AD dementia, and no significant interactions (Supplementary Table 2, 3).

**Fig. 2:**
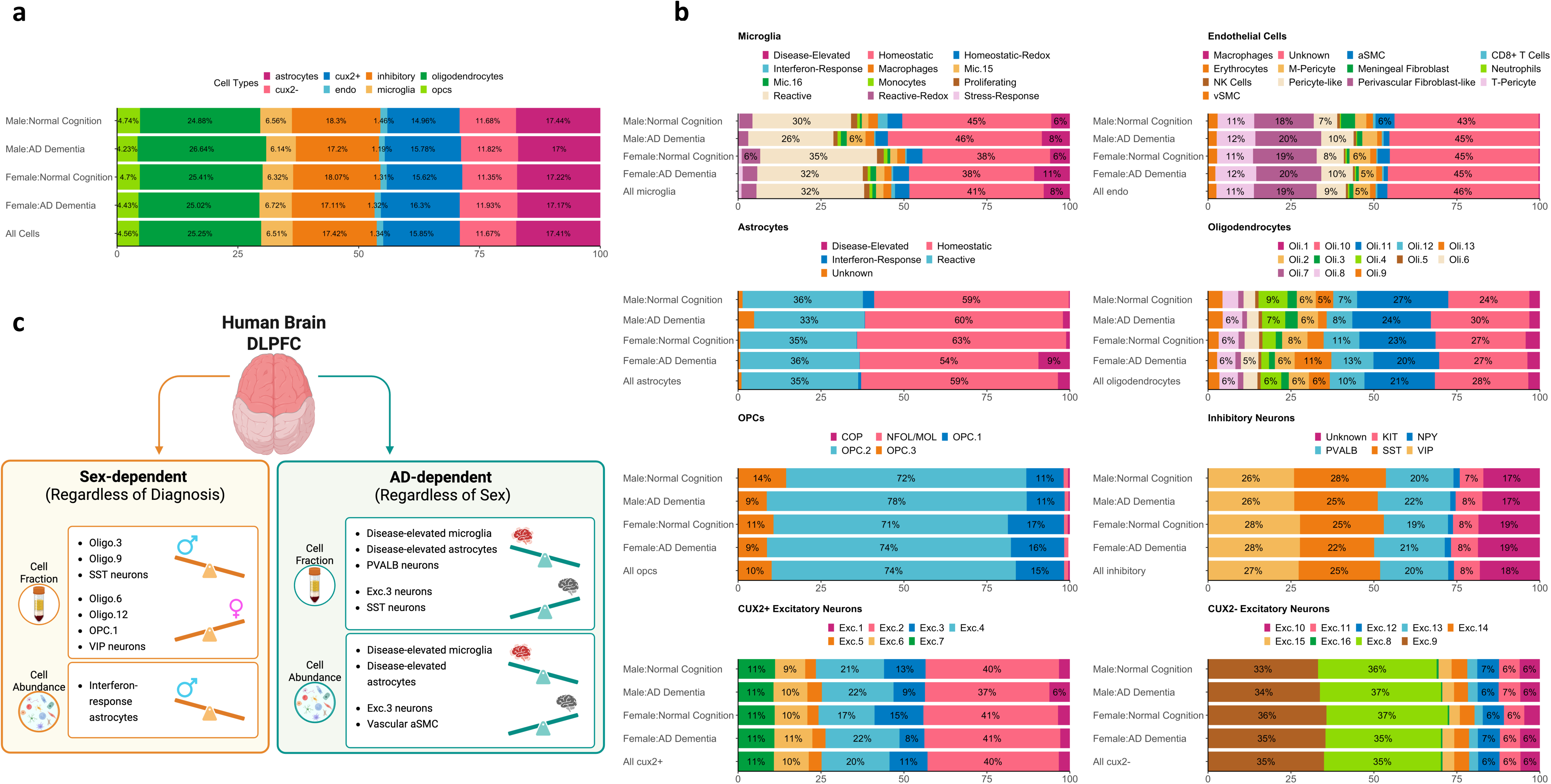
ROS/MAP snRNA-seq cell composition comparison. a, The proportion of each major cell type in ROS/MAP snRNA-seq samples by sex and diagnostic groups. Cell type abbreviation: cux2+/cux2-, CUX2+ or CUX2-excitatory neurons; inhibitory, inhibitory neurons; endo, endothelial cells. b, The proportion of each subtype of each major cell type in ROS/MAP snRNA-seq samples by sex and diagnostic groups. c, Sex-dependent or AD dementia diagnosis-dependent cell composition changes. Abbreviations: DLPFC, dorsolateral prefrontal cortex. Panel c. was created in BioRender.

Then, for the cell fractions of the 67 annotated cell subpopulations (Fig. 2b), seven showed a significant sex difference and four were significantly influenced by AD dementia. The cell types that showed the most difference in cell fractions either between sex or diagnostic groups were oligodendrocytes and inhibitory neurons. Females had a higher fraction of enhanced-translation Oli.6, Oli.12, enhanced-mitophagy subgroup OPC.1, and VIP inhibitory neurons, while males had a higher fraction of Oli.3, Oli.9, and SST inhibitory neurons (Fig. 2c, Extended Data Fig. 1). In AD, we observed higher fractions of disease-elevated microglia and astrocytes as expected, together with lower fractions of arterial smooth muscle cells (SMC.1 aSMC) within the vascular niche and excitatory neuron subtype 3 (Exc.3, mapped to cortical layer L2 and L3 by the Allen Brain Map; Fig. 2c, Extended Data Fig. 1). Notably, no significant sex and AD dementia diagnosis interaction effect was observed, suggesting cell type alterations in disease are largely shared across sexes (Supplementary Table 2).

From the cell abundance analysis, only interferon-response astrocytes showed a sex-dependent effect. Moreover, the same four cell subgroups showed an AD dementia-dependent effect as seen from cell fraction analysis (Fig. 2c). Specifically, males showed elevated interferon-response astrocytes compared to females (Extended Data Fig. 1), although the overall fraction of this cell subgroup is low in both sexes (2.4% for males, 0.1% for females). Increased disease-elevated microglia and astrocytes, along with decreased aSMC and Exc.3, associated with AD dementia (Extended Data Fig. 1). Again, no sex by AD dementia interaction effect was observed within any cell subpopulation (Supplementary Table 3).

### Overview of sex-specific associations with AD endophenotypes

Among the four AD endophenotypes studied, we identified 2,660 sex-specific transcript associations across eight major DLPFC cell types (Supplementary Table 4). Among the sex-specific associations, 79% were female-specific (likely due to sample size differences by sex), and about half (51%) were associated with Aβ (Fig. 3a). Neurons contributed to the most sex-specific associations, accounting for 62% of the total, whereas each glial cell type contributed less than 10% respectively (Fig.3b).

**Fig. 3:**
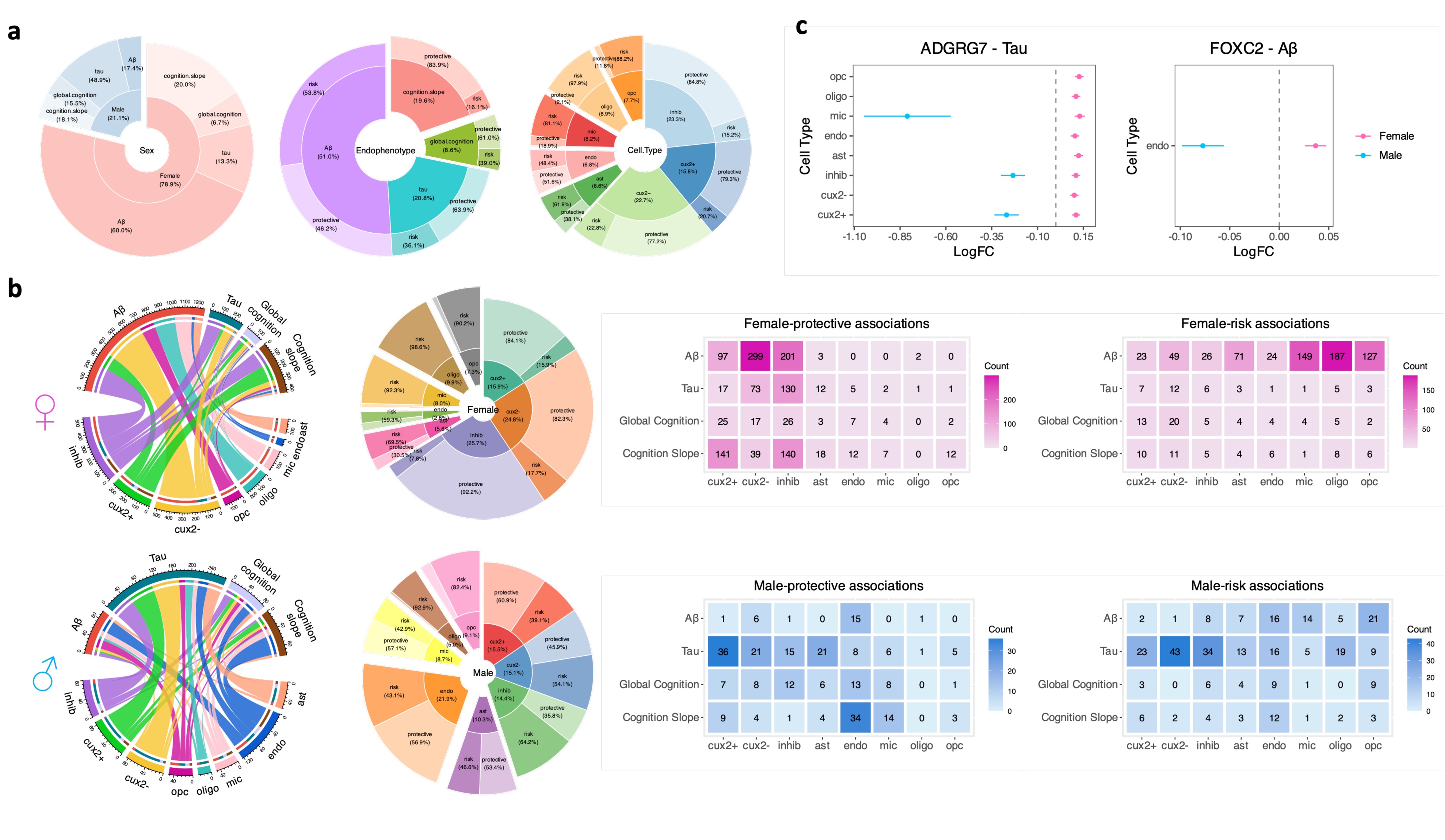
Sex-specific gene associations with AD endophenotypes. a, Pie-donut chart showing the relative proportions of sex-specific associations (N = 2,660) by sex, endophenotype, major cell types, and associations effect on four endophenotypes. Cell type abbreviations: ast, astrocytes; cux2+/cux2-, CUX2+ or CUX2-excitatory neurons; inhib, inhibitory neurons; mic, microglia; oligo, oligodendrocytes; opc, oligodendrocyte precursor cells; endo, endothelial cells. b, Comparison of female-specific vs male-specific associations distribution among eight major cell types and four endophenotypes by chord diagram, pie-donut chart, and heatmap. c, Forrest plot highlighting sex-specific associations in the same cell type with ADGRG7 and FOXC2 showing opposing effect between sexes for Tau and Aβ, respectively. Abbreviations: LogFC, log2 fold change.

Among the sex-specific gene associations, there were ten genes that showed FDR significant associations within each sex for the same AD endophenotype in the same cell type: *ADGRG7*, *EYA1*, and *FOXC2* for Aβ; *ADGRG7*, *ASIP*, *GPCPD1*, *KCNMB3*, *MRAS*, *SLC44A1*, and *UQCC1* for tau; and *IFI44L* for cognitive trajectory (Supplementary Table 5). Most of these associations displayed modest differences in effect size across sexes. However, interestingly two genes showed a risk effect in females but a protective effect in males, including *FOXC2* in endothelial cells on Aβ (risk for females, protection for males), and *ADGRG7* on tau from three cell types (*CUX2*+ excitatory neurons, inhibitory neurons, and microglia; Fig. 3c, Supplementary Table 5).

### Neurons may protect and glial cells may pose risk to A**β** in females

Among the female-specific associations we identified, 60% were with Aβ (Fig. 3b). In fact, the vast majority (77%) of detrimental gene associations among females were from glial cells in relation to Aβ (Fig. 3b). In contrast, the majority (86%) of neuronal associations in females demonstrated a protective effect, with 55% female protective associations across endophenotypes originating in excitatory neurons (Fig. 3b).

### Excitatory neurons may be the key cell type for tau associations for males

Among the male-specific associations we identified, 49% were with tau, while excitatory neurons (31%) and endothelial cells (22%) combined contributed to over half of all male-specific associations (Fig. 3b). Interestingly, neurons were responsible for providing the most protection against tau among males, especially CUX2+ excitatory neurons (Fig. 3b). In contrast to the female-specific effects outlined above, we did not see a striking split between neuronal and glial cell types with regards to detrimental and protective effects. Rather, male-specific associations showed more of a balance between detrimental and protective effects across cell types. Among all major cell types, endothelial cells seemed to play the most important role in protecting against increasing Aβ and faster cognitive decline among males (Fig. 3b).

### Validating sex-specific associations in SEA-AD cohort

To validate the 2,660 ROS/MAP sex-specific associations, we tested them in the independent SEA-AD cohort. To compare results from the two studies which used different cell taxonomies, we identified gene associations that showed a consistent effect with subgroups of the same parent cell type. We replicated nine female-specific associations involving eight unique genes, including *ADGRV1* and *OR3A3* with Aβ; *IFI27L1*, *LYRM1*, *STAP2*, and *TSTD2* with tau; *PDYN* with global cognition; and *TMEM50B* with cognitive trajectory (Fig. 4a). *ADGRV1*, *OR3A3*, and *PDYN* showed associations in excitatory neurons; *ADGRV1*, *TSTD2*, *IFI27L1*, *LYRM1*, and *STAP2* in inhibitory neurons; and *TMEM50B* in astrocytes. Higher *ADGRV1* and *PDYN* expression was related to worse AD endophenotypes, while elevations for the other genes were related to better AD endophenotypes (Fig. 4a). Interestingly, we also replicated the pattern of sex-specific effects in AD across cell types and endophenotypes, such that female-protective associations were enriched in neurons (93% for ROS/MAP, 80% for SEA-AD), whereas male-protective associations were evenly distributed across cell types (Fig. 4b). In addition, among all glial cell types, endothelial cells contributed to the most sex-specific associations in males (40% for ROS/MAP, 31% for SEA-AD; Fig. 4b).

**Fig. 4:**
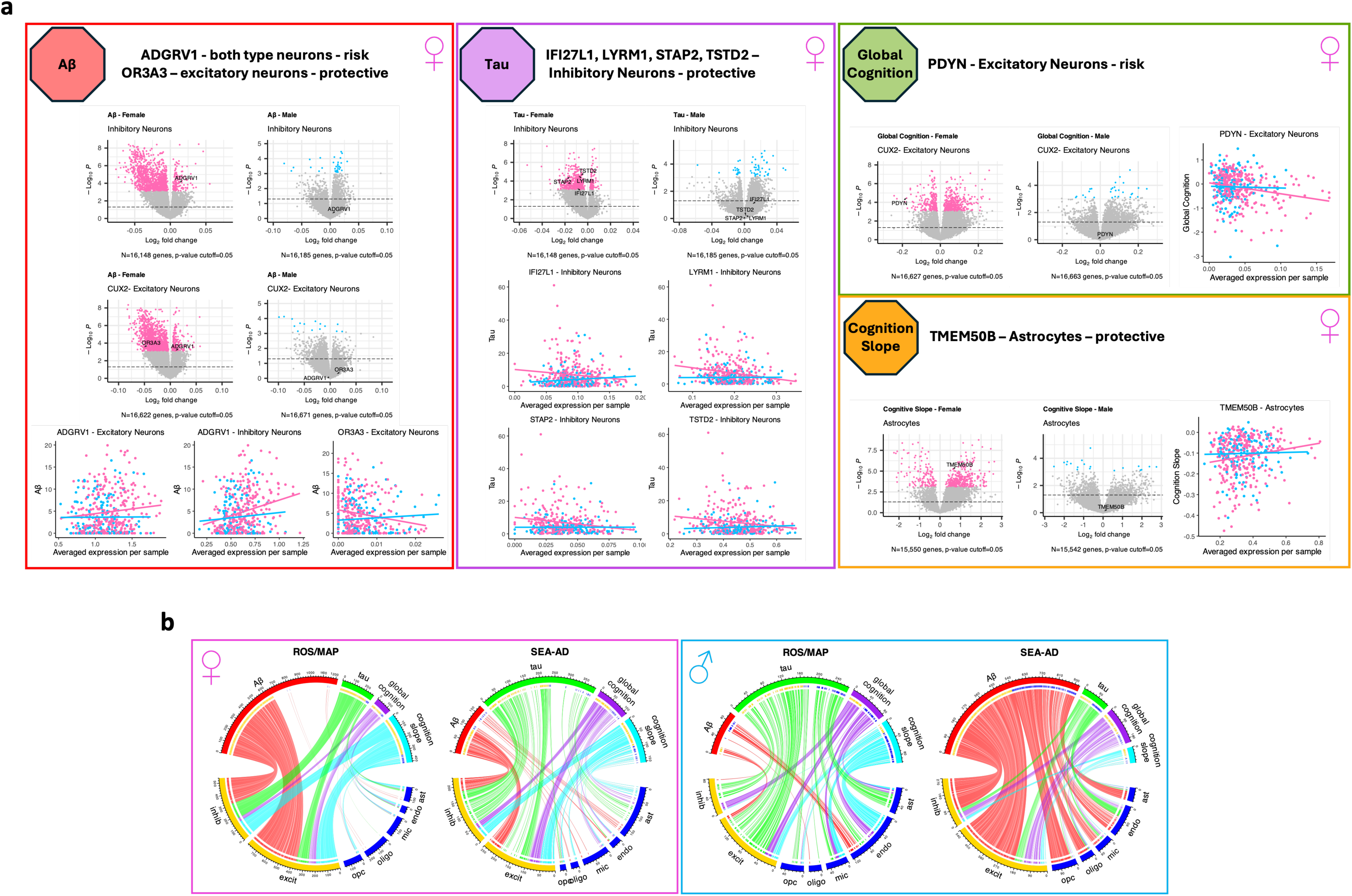
Replication of ROS/MAP sex-specific gene associations in SEA-AD. a, Replicated ROS/MAP sex-specific associations in SEA-AD cohort, all being female-specific associations. Volcano plot showing log2 fold change of gene expression associated with one unit increase of annotated endophenotype in each sex. Pink dots represent FDR associations in females; blue dots represent FDR associations in males; grey dots represent non-FDR results. Scatter plot showing correlation of highlighted gene expression and the annotated endophenotype in specific cell type. Pink dots represent female samples; blue dots represent male samples. Outliers beyond 4SD were removed from scatter plots. Lines in scatter plots represent the unadjusted linear fit. b, Chord diagrams showing sex-specific associations in females and males from ROS/MAP and SEA-AD cohorts. The length of each sector of the circle represents the total number of associations. Only protective links are displayed in the plots. Cell type abbreviations: ast, astrocytes; excit, excitatory neurons; inhib, inhibitory neurons; mic, microglia; oligo, oligodendrocytes; opc, oligodendrocyte precursor cells; endo, endothelial cells. Cell types were color-coded to show difference between neurons and glial cell types.

### Very few of the genes that show sex-specific associations with AD endophenotypes are differentially expressed by AD dementia diagnosis or by sex

We evaluated whether the sex-specific associations we observed are primarily driven by genes that are differentially expressed in disease, or genes that are differentially expressed by sex. Overall, we found 111 out of 2,472 (4%) sex-specific gene-cell type combinations showed differential expression patterns in AD dementia, involving 102 unique genes, 96 combinations in females, 15 in males (Fig. 5a, Supplementary Table 6). Similarly, 40 gene-cell type combinations from 31 unique genes (1%) were differentially expressed between males and females with AD dementia and only 10 gene-cell type combinations from 10 unique genes (0.5%) were differentially expressed between males and females among those with normal cognition (Fig. 5b, Supplementary Table 7). Male-specific effect genes from microglia showed the most (44%) differential expression pattern regardless of diagnosis, whereas female-specific effect genes from excitatory neurons had the most (45%) differential expressions. Three genes on sex chromosomes, *DDX3Y (*on Chr.Y), *TSIX* (on Chr.X), and *TXLNG* (on Chr.X), showed sex-biased expression (FDR<0.05) in both diagnostic groups, likely reflecting physiological differences by sex irrespective of disease for these genes. Due to the complexity of sex-dependent gene dose regulation on sex chromosomes, we will discuss genes on sex chromosomes further in a separate section below.

**Fig. 5:**
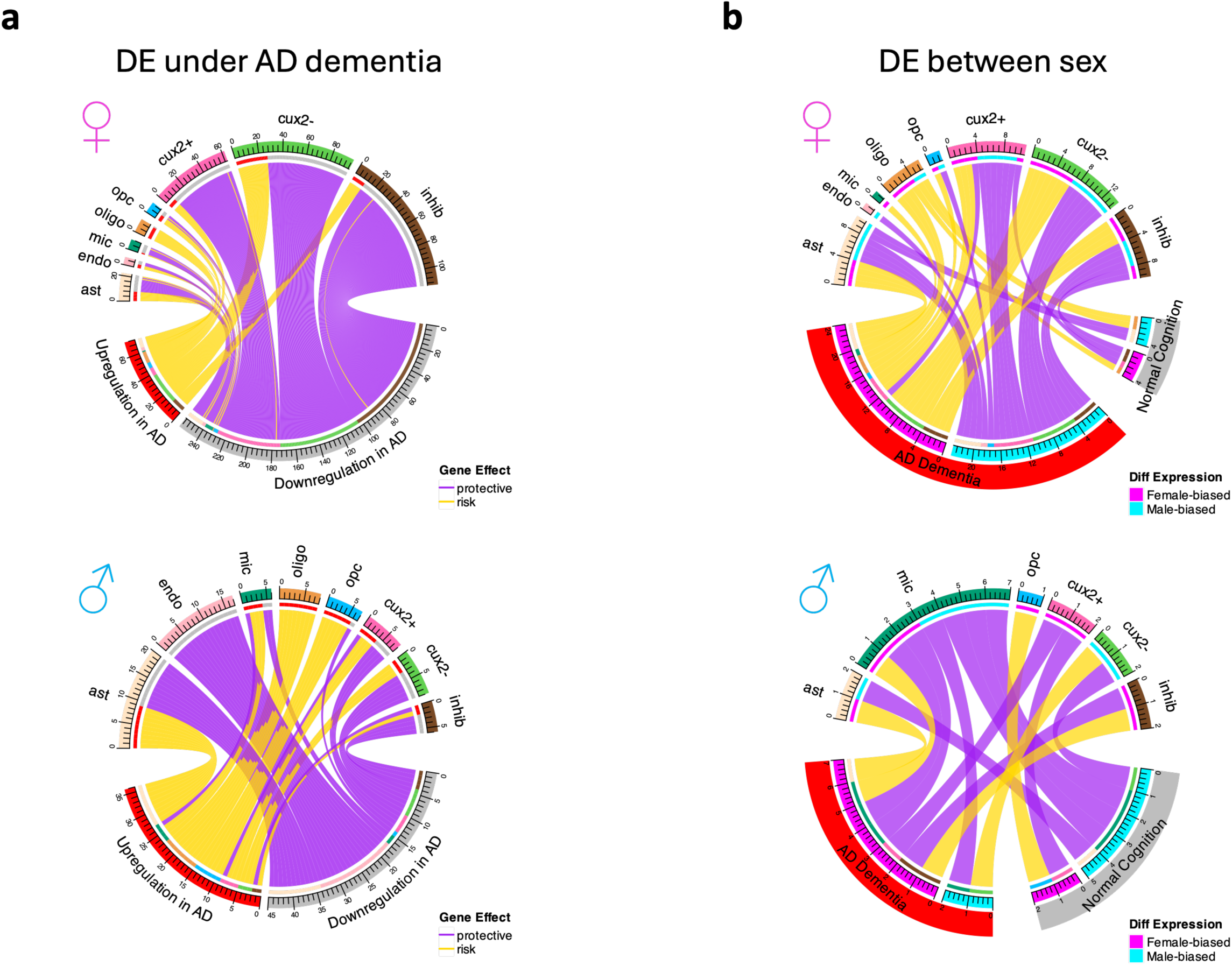
Differential expression of sex-specific association genes between sexes and under AD diagnosis. a, Chord diagram showing sex-specific association genes that FDR-differentially expressed pattern between sexes either in normal cognition or AD dementia group. b, Chord diagram showing sex-specific association genes that FDR-differentially expressed under AD dementia diagnosis. Purple links represent protective sex-specific associations; Yellow links represent risk sex-specific associations. DE, differential expression. Cell type abbreviations: ast, astrocytes; cux2+/cux2-, CUX2+ or CUX2-excitatory neurons; inhib, inhibitory neurons; mic, microglia; oligo, oligodendrocytes; opc, oligodendrocyte precursor cells; endo, endothelial cells.

### Sex-specific effect genes showed no over-representation on sex chromosomes

Among 2,110 sex-specific effect genes (the number of unique genes that make up the 2,660 sex-specific associations), 67 are on X chromosome and one is on Y chromosome, which is *DDX3Y* (Supplementary Table 4). We did not see an enrichment of sex-specific gene associations on the sex chromosomes relative to the autosomes either by total number (autosome vs. sex chromosomes: female-unique: 71 vs. 48, male-unique: 20 vs. 9, shared: 2 vs. 2; Fig. 6a) or ratio (autosome vs. sex chromosomes: female-unique: 4.8% vs. 3.4%, male-unique: 1.3% vs. 0.9%, shared: 0.2% vs. 0.1%; Fig. 6b). X chromosome genes are subject to X chromosome inactivation (XCI)^41^ in females, but due to being able to escape from XCI^42,43^ (a tissue-dependent process^44^), X-chromosome genes are enriched for having sex-biased expression^45–47^. Hence, to filter out this compounding factor for interpreting our result, we checked genes’ XCI patterns throughout our discovery process against a recent published XCI consensus call list^48^. For the 67 genes that show sex-specific effects on the X chromosome, 35 are subject to XCI and three are known XCI escapees, including *STS*, *SYAP1*, and *TXLNG* (Supplementary Table 4).

**Fig. 6:**
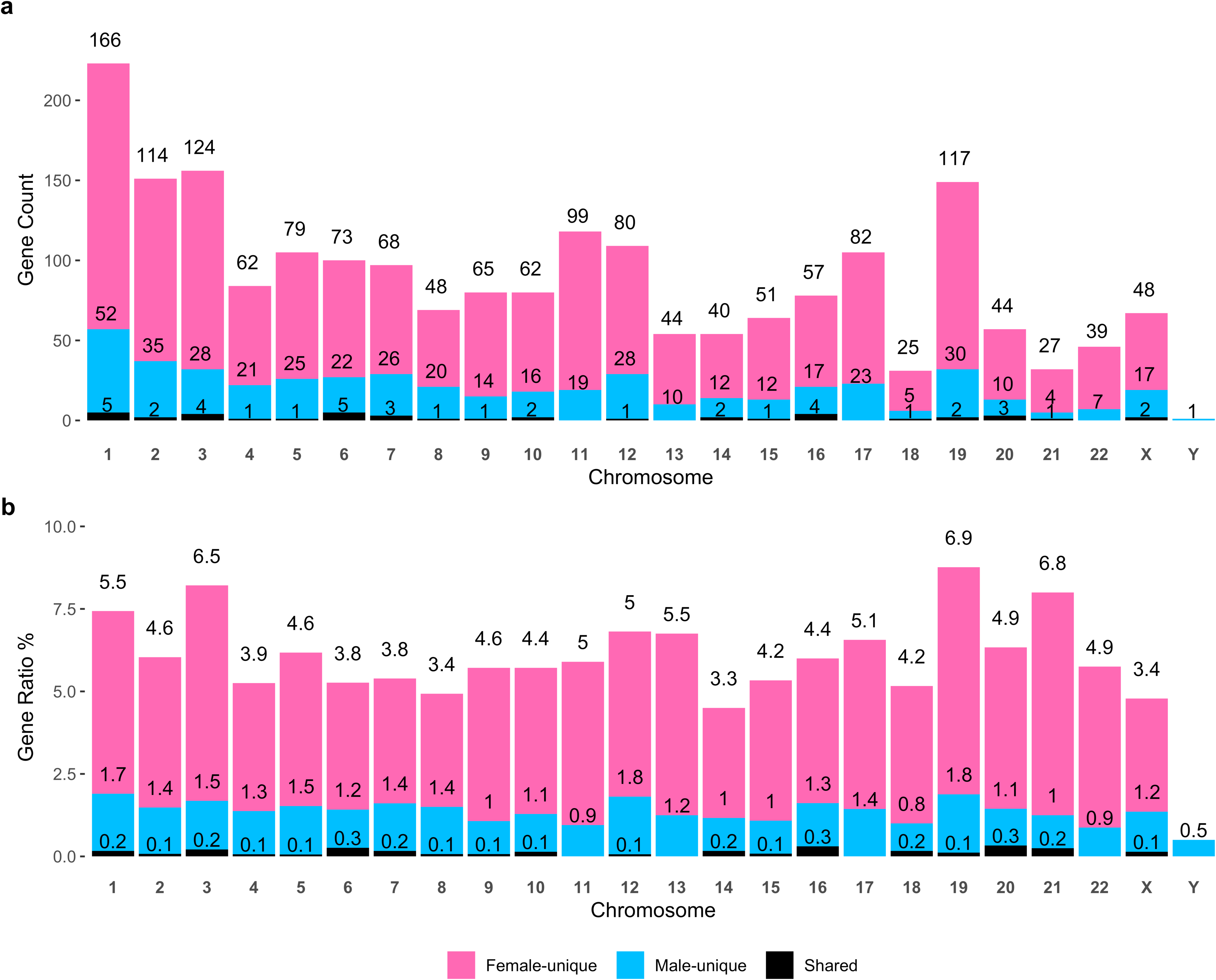
Chromosome localization of ROS/MAP sex-specific association genes. a, The count of sex-specific association genes on each chromosome. b, The ratio of sex-specific association gene count to total annotated gene number for each chromosome. Total annotated gene number of each chromosome was extracted based on Human Genome Assembly GRCh38.p14. Female-unique represents genes that only show sex-specific associations with females, male-unique represents genes that only show sex-specific associations with males, shared represents genes that show sex-specific associations with both sexes.

In addition, we more closely examined sex-biased expression patterns of these 68 genes from the sex chromosomes. Among these genes, seven showed sex-biased expression in the cell type they exerted sex-specific effect under either normal cognition or AD dementia, including *BX842570.1* (inhibitory neuron), *DDX3Y* (microglia), *STS* (CUX2+ excitatory neuron), *TSIX* (CUX2+ excitatory neuron), *TXLNG* (oligodendrocyte), *ASMTL-AS1* (CUX2-excitatory neuron), and *SYAP1* (oligodendrocytes). *DDX3Y* is the only Y chromosomal gene from our list, and as expected, it showed male-biased expression regardless of disease status.

### Eight sex-specific effect genes were known AD risk genes from GWAS studies

AD GWAS studies to date have identified over 100 genomic loci to be linked to AD disease susceptibility^49^. We examined our sex-specific effect gene list against 112 recent-reviewed AD risk loci from GWAS^49^, and found eight GWAS associated genes that are on our list, including *CR1*, *WDR12*, *INPP5D*, *HESX1*, *TNIP1*, *EPHA1*, *SHARPIN*, and *CTSH.* All these genes showed sex-specific associations with either Aβ or tau pathology. *CR1* showed a male-specific effect, while the rest showed female-specific effects (Supplementary Table 4). Higher levels of *CR1* in microglia were associated with less tau among males. For the female-specific associations, higher levels of five genes in neurons were associated with better AD endophenotypes: including *EPHA1, HESX1,* and *WDR12* in CUX2-neurons with Aβ, and *SHARPIN* in both CUX2- and inhibitory neurons with tau. In contrast, among the glial cells, higher *INPP5D* in astrocytes and *CTSH* and *TNIP1* in microglial was related to a higher Aβ. While sex-specific associations for the lead variants at these candidate genes have not been reported in the literature.

### Sex-stratified association genes are enriched for immune/inflammatory/stress response pathways

In gene-set enrichment analyses, we observed many sex-specific pathways (Fig. 7a, Supplementary Table 8). For females, gene associations with cognitive performance contributed to 66% of female-specific enrichment pathways, whereas for males, enriched pathways were distributed evenly among the four AD endophenotypes. Male-specific pathways included “ANGIOGENESIS”, “APOPTOSIS”, “COMPLEMENT”, “ESTROGEN_RESPONSE_EARLY”, “IL2_STAT5_SIGNALING”, “INFLAMMATORY_RESPONSE”, “KRAS_SIGNALING_UP”, and “TGF_BETA_SIGNALING”. Female-specific pathways included “APICAL_JUNCTION”, “BILE_ACID_METABOLISM”, and “XENOBIOTIC_METABOLISM”.

**Fig. 7:**
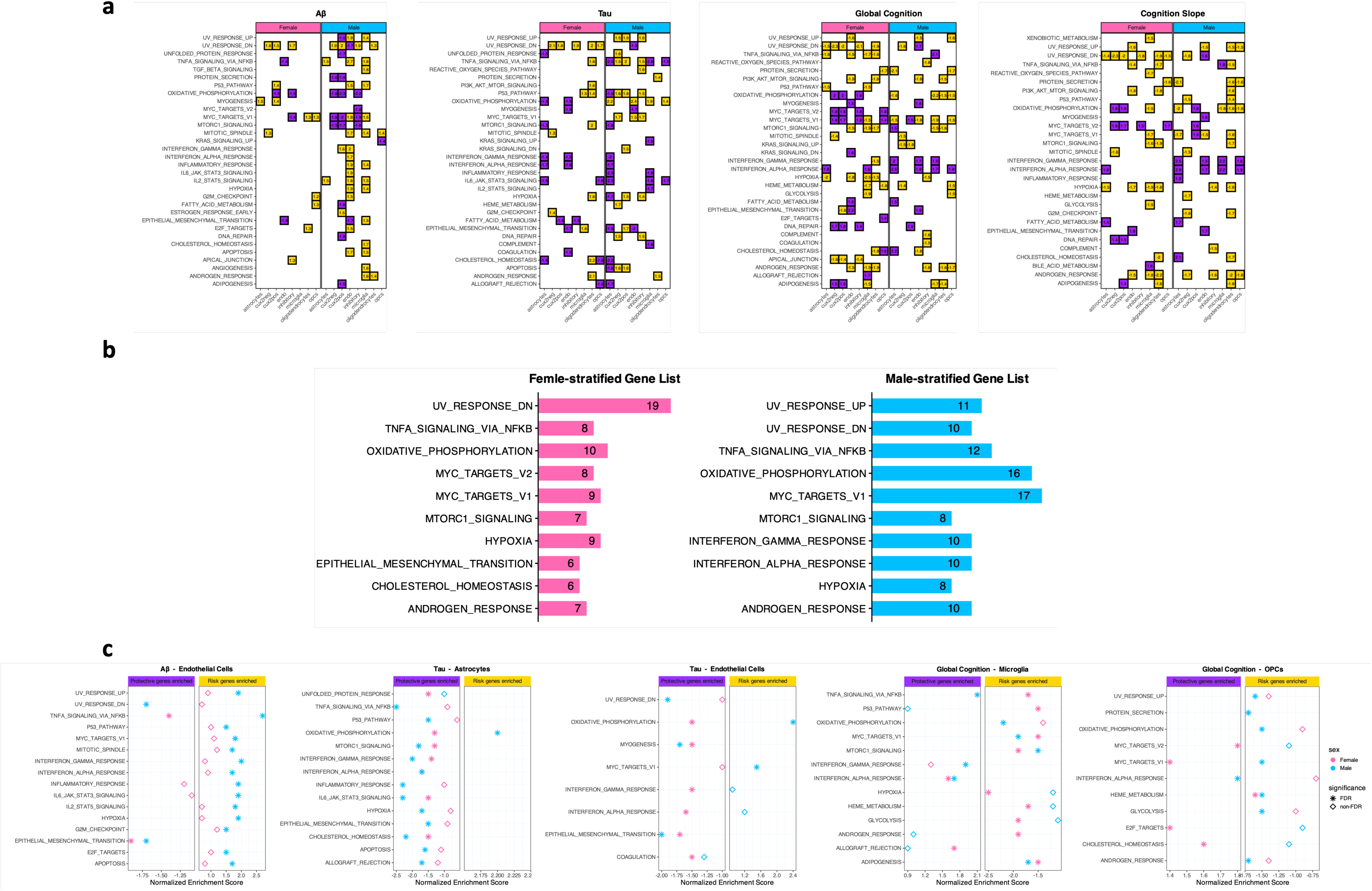
Gene set enrichment pattern of sex-stratified AD endophenotype associations. a, ALL global false discovery rate (FDR) <0.05 gene set enrichment results from sex-stratified associations for each of four endophenotypes tested. Purple tiles indicate risk association genes enriched; yellow tiles indicate protective association genes enriched. Annotated numbers are normalized enrichment scores extracted from the analysis. b, Top 10 FDR enrichment pathway by number for each sex. Annotated numbers showing the total frequency of individual pathway was seen FDR-enriched. c, Five combinations of endophenotype and cell type that showed enrichment of the same pathway with opposing effect gene enriched between sex.

Interestingly, we observed three pathways that were enriched in opposing directions by sex (Fig. 7c). The first one was “TNFA_SIGNALING_VIA_NFKB” which was enriched in opposite directions for microglial gene associations with cognition and endothelial gene associations with Aβ. In both cases, the pathway was enriched for protective gene associations among males, but detrimental gene associations among females. The second pathway showing this pattern is “OXIDATIVE_PHOSPHORYLATION” in both astrocytes and endothelial cells, in this case with protective gene associations among females and detrimental gene associations among males. Finally, “MYC_TARGETS_V1” in OPCs was enriched for protective gene associations with cognition among females, but detrimental gene associations with cognition among males.

Regardless of differences, both sexes shared many enriched pathways (Fig. 7b). There were 22 unique pathways across all major cell types and endophenotypes that were enriched in both sexes in the same cell type for the same AD endophenotype in same direction (Supplementary Table 8), accounting for one third of all female enrichments and a quarter of all male enrichments. These common pathways highlight the shared molecular mechanisms of AD pathogenesis between sexes as expected.

### Sex-specific effects led to sex-differences in intercellular communication in the AD dementia brain

To get an overview of sex differences in cell-cell communication patterns, we first inferred intercellular communication of all curated signals in each sex and diagnostic group. All four groups had comparable communication count and strength (probability), except for a significant reduction (adjusted p-value < 0.05) of communication strength among AD females compared to normal cognition males (Fig. 8a). The strongest intercellular communications among DLPFC cell types were among neurons, astrocytes, and OPCs for all groups (Fig. 8b). Among those with normal cognition, females showed a slightly stronger (p>0.05) communication strength from OPCs to oligodendrocytes than males. However, the difference disappeared among participants with AD (Fig. 8c), due to the reduction of such communication pathways in females. On the other hand, AD females showed a slightly stronger (p>0.05) communication strength within microglia compared to AD males (Fig. 8c), due to a larger downregulation of such communication pathways in males with AD. For both sexes, the biggest reduction in cell communication among those AD was observed among excitatory neurons, inhibitory neurons, and OPCs (Fig. 8d). On an individual communication signal level, we observed several that were differently dysregulated in AD between sexes that did not present among those with normal cognition (Fig. 8e). For example, compared to AD females, AD males lacked meaningful inferred communication of the THY1, COMPLEMENT, MHC-II, and ICAM signals, due to significant downregulation of these pathways among those with AD dementia. Interestingly, all of these are exclusively incoming signals from microglia that were present in males with normal cognition, except for THY1 sent from endothelial cells (Fig. 8f), which was unique to females. Conversely, AD females lacked significant communication of CD99, due to a significant reduction in this cell communication pathway among females with AD dementia (Fig. 8f). CD99 showed an outgoing signal from both endothelial cells and astrocytes in males regardless of diagnosis; however, females with normal cognition only showed CD99 sent from endothelial cells (Fig. 8f), indicating astrocyte-sent CD99 signal is unique to males.

**Fig. 8:**
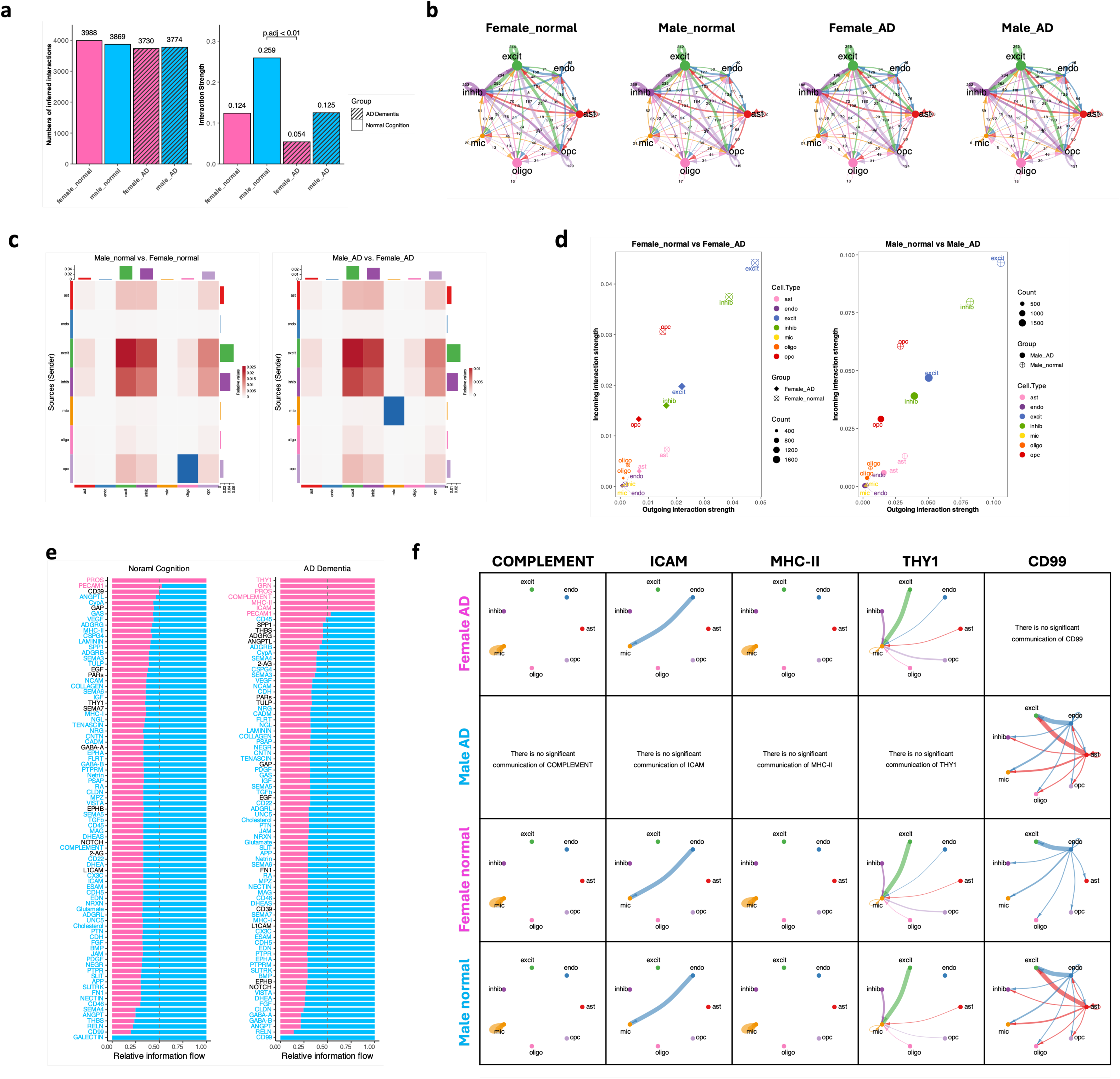
Intercellular communication comparison between sex and AD dementia diagnostic groups. a, Comparison of total count and strength of all curated intercellular communication interactions for four groups. b, Overall information flow of four groups. Annotated numbers showing the total count of interactions between each cell type pair. Cell type abbreviations: ast, astrocytes; excit, excitatory neurons; inhib, inhibitory neurons; mic, microglia; oligo, oligodendrocytes; opc, oligodendrocyte precursor cells; endo, endothelial cells. c, Differential interaction strength heatmap between sexes among those with normal cognition or AD dementia. The colored bar plot at the top represents the sum of incoming signaling displayed in the heatmap. The colored bar plot on the right represents the sum of outgoing signaling. The bar height indicates the degree of change in terms of the number of interactions or interaction strength between sex. In the heatmap, red (or blue) represents increased (or decreased) signaling in the male dataset compared to the female dataset. d, Changes of the outgoing and incoming interaction strength for each major cell type in each sex with or without AD dementia. e, Comparison the information flow between sexes for each signaling pathway among each diagnostic group. Each inferred signaling network is defined by the sum of the communication probability (strength) among all pairs of cell groups in the inferred network. Significant signaling pathways were ranked based on differences in the overall information flow within the inferred networks between sexes. A paired Wilcoxon test is performed to determine whether there is a significant difference in the signaling information flow. Signaling pathways colored pink are enriched in females; those colored blue are enriched in males; and those colored black show no difference between sexes. f, Circle plot of intercellular communications of COMPLEMENT, ICAM, MHC-II, THY1, and CD99 networks in four groups. Circle sizes representing major cell type are proportional to the number of cells in each cell group. The width of chords represents the strength of the total communication.

We then identified unique outgoing or incoming signals from each of the major cell types between sexes and diagnostic groups in a pairwise fashion. We found a total of 124 unique incoming or outgoing signals which represented 34 unique pathways among the four groups (Supplementary Table 9). Almost one-third (30%) of these unique signals involved microglia, hinting at its leading role in driving sex differences in intercellular communication networks in AD. AD females (32%) and normal cognition males (33%) combined to contribute two-thirds of all unique incoming or outgoing signals (Supplementary Table 9), indicating downregulation of male-specific intercellular communications in AD males and upregulation of AD-specific signals in AD females as the main mechanisms underlying sex-specific cell signaling alterations in AD.

Next, we examined the involvement of the 2,110 sex-specific effect genes in these 34 unique communication networks. Only 133 of our sex-specific effect genes were curated in human CellChatDB v2, and they are involved in 119 intercellular signals by encoding either a ligand or a receptor. Of these, six pathways including three sex-specific effect genes (*ITGB1*, *JAM2*, and *TNC*) showed communication alterations that aligned with the direction of effect and the cell type implicated in our sex-specific gene association analyses above. These signals are COLLAGEN, FN1, JAM, LAMININ, SPP1, and TENASCIN, all of which contribute to unique incoming signals to microglia among females with AD dementia compared to males with AD, and among males with normal cognition compared to males with AD dementia (Supplementary Table 9). Though it should be noted that each of these signals includes numerous cell types besides microglia (Extended Data Fig. 2). Upon further investigation, these signals share a common receptor on microglia, ITGB1 (encoded by *ITGB1*). In all cases, the sex difference in this signaling pathway was driven by a more severe downregulation of incoming ITGB1 signals to microglia among males with AD to the degrees that there were no inferred communications of these signals in this cohort (Fig. 9a). Microglial *ITGB1* indeed showed more expression in females with AD compared to males with AD (p=0.02, FDR=0.4; Supplementary Table 7), but no sex-by-AD interaction effect was observed (p=0.06). Notably, *ITGB1* in microglia shows a risk association with Aβ among females (FDR=0.01, Fig. 9b), suggesting these six microglial incoming signals might play a role in sex-specific risk for Aβ accumulation.

**Fig. 9:**
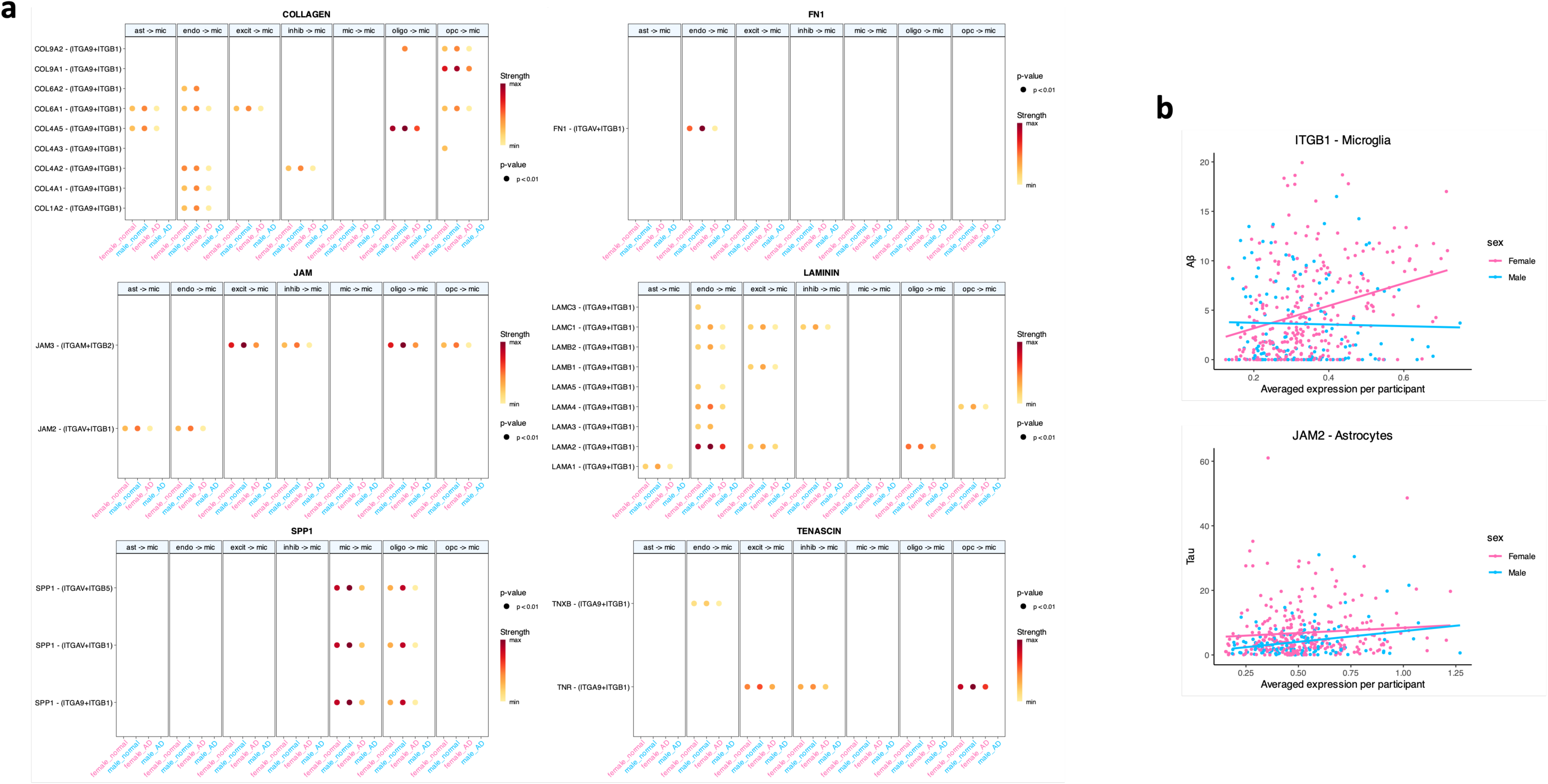
ITGB1 and JAM2 induced six sex-unique microglial-incoming signals under AD dementia. a, Dot plot showing all enriched intercellular communication (L-R pair interactions) from every major cell group to microglia, evaluated by sex and diagnosis. b, Scatter plot showing the correlation of ITGB1 expression with Aβ in microglia and JAM2 expression with tau in astrocytes. Pink dots represent female samples; blue dots represent male samples. Outliers beyond 4SD were removed from scatter plots. Lines in scatter plots represent the unadjusted linear fit.

Apart from using ITGB1 as a receptor, the JAM signal also involves another sex-specific effect gene *JAM2*, which encodes one of its ligands by the same name (JAM2). The JAM2-(ITGAV+ITGB1) interaction induced a unique incoming signal to microglia from astrocytes and endothelial cells in AD females (Fig. 9a). *JAM2* expression in astrocytes was significantly increased (FDR=0.03) in AD dementia compared to normal cognition in both sexes (female p= 0.004, male p=0.09), but it did not show sex-biased expression in any cell type regardless of disease diagnosis. Hence, the lack of JAM2-(ITGAV+ITGB1) signal to microglial in AD males was due to the downregulation of *ITGB1* as we discussed above. Since *JAM2* is a male-specific risk gene in astrocytes for tau (FDR=0.02, Fig. 9b), we hypothesize JAM2 induced JAM signal from astrocyte to microglia functions as a protective mechanism against tau pathology for men. Lastly, TNC is curated to be a ligand for TENASCIN signals; however, no meaningful communications induced by it was observed in any of the four groups we analyzed.

## DISCUSSION

We have provided a comprehensive assessment of sex-specific transcriptomic effects that contribute to AD neuropathology and cognitive decline in a cell-specific manner (Fig. 10a). Our results highlight a striking dissociation between protective and detrimental gene associations among females, with protective associations arising primarily from neurons and detrimental gene associations arising primarily from glial cells. In contrast, we did not observe a similar dissociation among males (Fig. 10b). We also identified and replicated nine cell-specific gene associations with AD neuropathology and cognitive decline, all of which act in a female-specific manner, highlighting a set of genes that may have therapeutic potential in women. Next, we identified a host of sex-specific biological pathways and cell-cell communication alterations that behave in a cell-specific manner (Fig. 10b), drawing attention to novel biological signals that may only be relevant among males or females. Among those sex-specific effects, we highlight *ITGB1* as a high-quality candidate for a sex-biased role in Aβ accumulation in AD that deserves further functional validation. Finally, in contrast to some work, we did not observe a substantial enrichment for sex-specific gene associations with AD endophenotypes on the sex chromosomes, suggesting that sex-biased expression on the X and Y chromosomes may not be a major contributor to sex-specific risk and resilience in AD.

**Fig 10:**
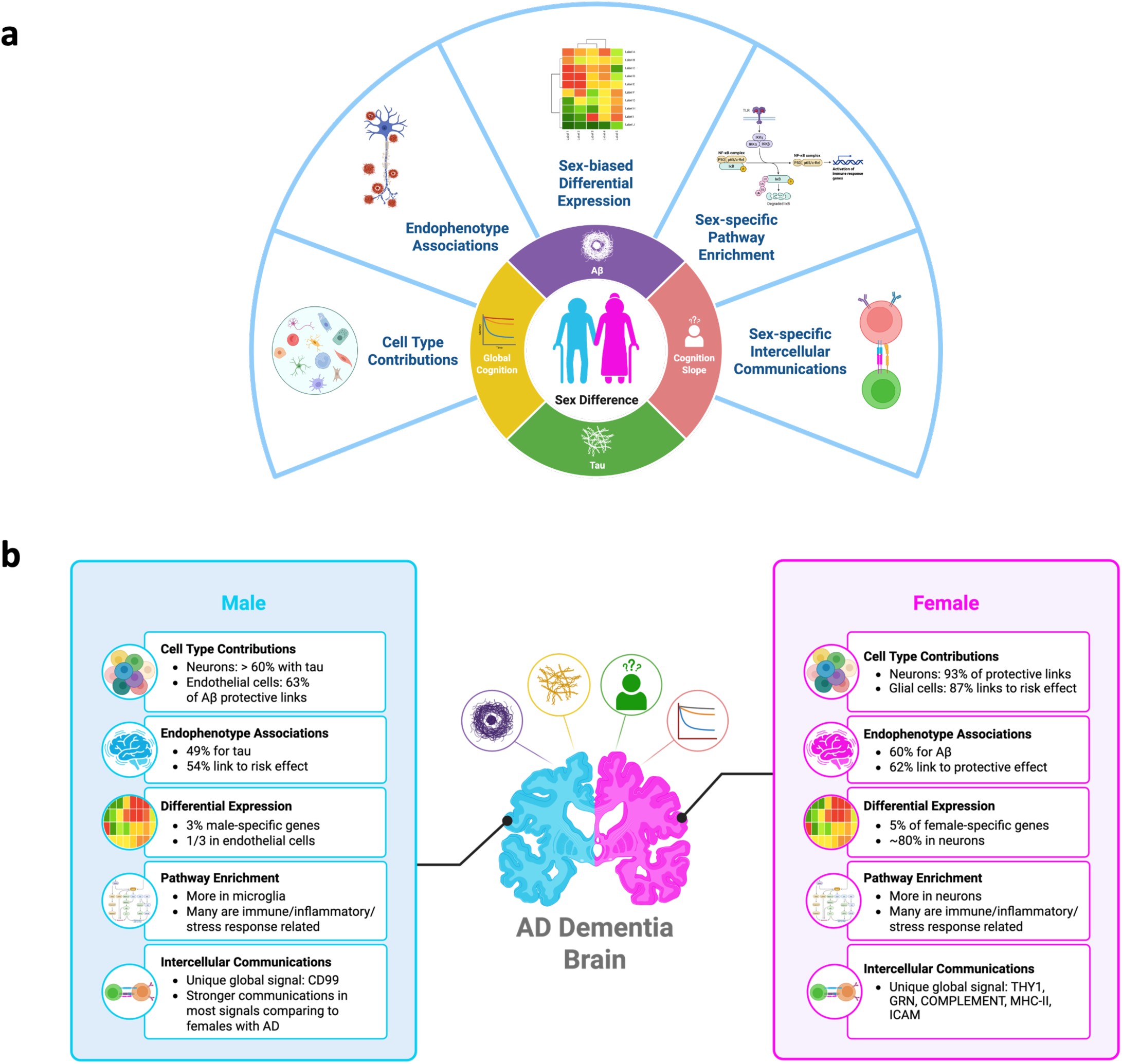
Summary of single cell landscape of sex-specific differences in AD dementia. a, Schematic illustration of the analysis strategy. b, Key sex-specific associations with AD endophenotypes at single cell resolution. Both figures were created in BioRender.

### A host of neuronal genes protect against AD neuropathology among females, while endothelial cell genes protect against A**β** and cognitive decline among males

Nearly all protective gene associations with AD neuropathology among females were observed in neurons (93%), suggesting a female-specific alterations in neurons that may be critical for understanding neuroprotection in AD. Our observation from these two independent cohorts that females show a sharper downregulation of protective genes in neurons seems to align with earlier studies in fewer participants showing a similar downregulation of gene expression in both excitatory and inhibitory neurons among females in response to AD pathology, a change that was not seen among males^50^. More interestingly, that same study also showed that the extreme changes observed in neuronal cells in females showed particularly strong associations with Aβ, aligning with our findings in the present study in which 50% of all protective neuronal associations in females were in relation to Aβ.

The female-specific associations between neuronal gene expression and Aβ aligns well with previous literature in other domains. Our group and others have shown that Aβ positive females are more susceptible to hippocampal atrophy and cognitive decline^51^, a finding that has been well-supported by the autopsy literature as well^4^. In fact, a similar neuronal vulnerability is also observed in mouse models of brain amyloidosis where female mice show faster neurodegeneration and cognitive decline in response to Aβ^52^. While response to Aβ pathology is the simplest interpretation and aligns well with the literature, we cannot rule out the possibility that these alterations are upstream of Aβ in females and protect against deposition. Future mechanistic work is needed to clarify the temporal ordering of the associations and clarify whether down regulation of neuronal genes in women is a cause or consequence of Aβ deposition.

In contrast to these female-specific effects, we observed robust male-specific protective associations in endothelial cells, particularly in relation to Aβ and cognitive decline. Endothelial cells made up the majority of male-specific protective associations with Aβ and cognitive decline, and also presented with the most enriched pathways among males. It is known that sex hormones influence endothelial cell health and function, and hence there are notable differences in endothelial cell structure and function between the sexes^53^. Endothelial cells have a sex-dependent secretome^54^ and 14-25% of the endothelial transcriptome is sex-biased, due to differences both at birth and acquired over life^55^. Our results suggest that these notable differences in endothelial cell function may lead to differential disease-response in the AD brain as AD neuropathology begins to accumulate.

Additionally, and in contrast to the female-specific associations highlighted above, male-specific associations in neurons were primarily in relation to tau. While females appear to accumulate tau pathology more rapidly than males^56,57^, and show a faster rate of cognitive decline in the presence of AD pathology as highlighted above including in relation to tau pathology^58^, there is some evidence it takes a higher level of tau pathology for females to reach a clinical diagnosis of AD^59^. Perhaps the male-specific and female-specific gene associations that we have observed in the present analysis provide some additional insight into these contradictory interpretations of similar data. It may be that the downregulation of protective gene effects in neurons among females arise more sharply in relation to Aβ and drive a faster rate of tau accumulation and a faster rate of cognitive decline. At the same time, males may show a stronger neuronal response to tau deposition that makes them more susceptible to the downstream consequences of tau.

From the present results, it is difficult to speculate on the direction or timing of these sex-specific effects given the cross-sectional nature of the study, but these results offer additional information about the clear sex differences in the cellular profile of the AD brain and highlight the clear need to better characterize how and when sex-specific associations emerge across stages of disease.

### Novel female-specific gene associations with AD neuropathology and cognitive decline

All our replicated sex-specific associations were for females, likely due to the larger sample size and additional statistical power. First, we observed three female-specific associations in neurons with Aβ including *ADGRV1* in both excitatory and inhibitory neurons and *OR3A3* in excitatory neurons. Both *ADGRV1* and *OR3A3* encode proteins that are members of G protein-coupled receptors, which mediate most of our physiological responses to hormones, neurotransmitters and environmental stimulants^60^. *ADGRV1* plays an important role in brain development and sensory cell function^61^. Mutations in *ADGRV1* have been linked to Usher syndrome type 2C (USH2C, OMIM 605472)^62^, the most common form of hereditary deaf blindness, and various forms of epilepsy^63,64^. USH2C can affect both sexes, although some reported males with *ADGRV1* mutations show more severe and even lethal clinical manifestations^65^. Additionally, one study identified *ADGRV1* as a male-specific risk factor in opioid dependence in African Americans^66^, highlighting potential differences in *ADGRV1* function by sex. *OR3A3* encodes an olfactory receptor protein, but its function in human disease is not well-defined. Certainly, there are notable impairments in olfaction in AD, so alterations in this gene may reflect sex differences in neurodegenerative process relevant to olfaction, aligning with recent evidence that females show a faster rate of neuronal loss in the olfactory cortex during AD compared to males^67^, and that there is a tighter coupling between olfaction impairment and cognitive impairment among females compared to males^68^.

We also observed four female-specific associations in neurons with tau including *IFI27L1*, *LYRM1*, *STAP2*, and *TSTD2*, all in inhibitory neurons. *IFI27L1* is part of the interferon-stimulated gene (ISG) family, which is typically involved in the immune response to viral infections and other inflammatory processes. IFI27 was identified as one of three ISG genes that mediated an antiviral effect in cortical neurons^69^; however, the particular evidence for IFI27L1 in human cells is still lacking. LYRM1 has been mostly studied for its role in regulating adipose tissue homeostasis and obesity-associated insulin resistance^70^. Overexpression of LYRM1 in adipocytes has been linked to mitochondrial impairment^71^. STAP2 was shown to be an adaptor protein to enhance TCR signaling^72^, dysregulation of which can either cause immunodeficiency or autoimmunity linked to diseases such as multiple sclerosis^73^. Studies of TSTD2 are limited, but serum TSTD2 antibody levels were reported to be higher in males, older adults, smokers, in those who consumed alcohol regularly, in those with hypertension, and significantly higher in patients with acute cerebral infarction or chronic kidney disease^74^.

Finally, we observed two female-specific associations in neurons with cognitive decline including *PDYN* in excitatory neurons and *TMEM50B* in astrocytes. *PDYN* encodes the dynorphin neuropeptides, polymorphisms and structural variants within this gene locus were reported to be associated with drug addiction^75^. Interestingly, many *PDYN* variants exclusively showed associations with risk in female opioid addicts^76^. *TMEM50B* encodes a membrane protein which presents on endoplasmic reticulum and Golgi apparatus. Located on chromosome 21, *TMEM50B* has been implicated in the neurophenotypes of Down Syndrome due to being one of a few genes upregulated in model systems of Down Syndrome, particularly in the cerebellum during development^77^.

### Immune/inflammatory response and microglia may play central role to sex differences in AD

Another key finding from our results is that immune and inflammation response pathways show striking sex-specific enrichment for associations with AD neuropathology, particularly in microglia, endothelial cells, and OPCs. Of particular note, the TNF-α signaling via NF-κB pathway and MYC Targets pathway were enriched in each sex with an opposite direction of association. The role of NF-κB signaling in AD pathogenesis, especially in glial cells, is well-known to produce pro-inflammatory cytokines and induce neuroinflammation, promote myelin injury and neurodegeneration, drive Aβ formation and tau hyperphosphorylation, and stimulate more NF-κB activation in neurons and glial cells^78,79^. The fact that NF-κB signaling showed sex-dimorphic enrichment might indicate a drastic sex difference in the microenvironment for these cell types during AD. In fact, our observations of many unique microglia-based signals in each sex seems to provide further support to this hypothesis.

Indeed, the importance of microglia-specific signals in contributing to sex differences in AD was evident from our intercellular communication analysis. We not only showed that microglia present with the most sex-specific enriched pathways but also showed that 30% of group specific intercellular signals involved microglia, significantly more than any other cell type. Microglia are the primary resident immune cells of the brain and play a crucial role in regulating the immune response to AD^80–82^. Many of the top AD genes identified in GWAS studies are expressed by microglia, such as *TREM2*, *ABCA7*, *CR1* and *INPP5D*. Moreover, increasing evidence has recognized the importance of microglia in contributing to sex-differences in AD^1^. Our intercellular communication profiling pinpointed several pathways that are differentially dysregulated in AD between sexes, including THY1, COMPLEMENT, MHC-II, ICAM, and CD99, many of which are well known for their roles in immune and inflammatory response.

It is also worth highlighting that one key player from microglia induced numerous sex-unique signals: *ITGB1,* a gene that encodes the β1 subunit of integrin receptors. *ITGB1* was identified as one of the five hub genes that is associated with the progression of AD and Major Depressive Disorder^84^. Most interestingly, microglial ITGB1 is a key component of the cell surface receptor system that mediate interaction between fibrillar Aβ (fAβ) and microglia to promote the clearance and phagocytosis of fAβ^85^. fAβ can induce overexpression of *ITGB1* transcription, whereas the more potent neurotoxicity form Aβ oligomers can downregulate *ITGB1*^86^. Our evidence that microglial *ITGB1* may act as a female-specific risk factor for Aβ suggests that there is a higher sensitivity of *ITGB1* to Aβ pathology in the female brain.

### Sex chromosomes are not enriched for sex-specific associations with AD endophenotypes

While there is emerging evidence of sex-biased expression of genes on the sex chromosomes, particularly the X chromosome, contributing to AD pathogenesis^83,87,88^, our study found that only 3.2% of all sex-specific gene associations were located on the X chromosome, accounting for less than 5% of all X-linked genes and indicating no enrichment on the X chromosome compared to the autosomes. Similarly, we only observed one sex-specific gene association on Y chromosome. Interestingly though, we did observe Androgen Response as one of the top pathways that was significantly enriched in multiple cell types for both sexes, highlighting the possibility of a sex-by-hormone interaction effect in AD.

Among the X-chromosome genes contributing to sex-specific associations we observed, a few are known XCI escapees. Oligodendrocyte-*TXLNG* showed female-biased expression regardless of disease, which aligns with the fact that it is a known XCI escapee in the brain^89^. *TSIX* from CUX2+ neurons, the antisense regulator of XIST, also showed female-biased expression in both diagnostic groups. Our observation is consistent with a previous study showing both *TXLNG* and *TSIX* had female-biased expression in human blastocysts from sibling embryos^90^, indicating a baseline expression difference between sexes that is independent of AD dementia. *BX842570.1* from inhibitory neurons showed female-biased expression only among participants with normal cognition (female-more expression in AD dementia with a FDR=0.07), but its XCI pattern is unknown, although it locates adjacent to a known and well-studied XCI escape gene *PRKX*^91^, about 100kb away on the opposite strand. *STS* from CUX2+ neurons, which encodes an endoplasmic reticulum protein called Steroid Sulfatase, and showed female-biased expression among participants with normal cognition. Since it is a known XCI escapee in the brain^48,89^, this expression pattern can be expected. Interestingly, *STS*’s sex-biased expression was not observed among participants with AD dementia. Oligodendrocyte-*SYAP1*, which encodes the Synapse Associated Protein 1, showed female-biased expression among participants with AD dementia. It is a known XCI escapee, and hence one would expect to see a female-biased expression pattern regardless of diagnosis. It is possible that the heterogeneity in sex-biased expression among those with AD dementia compared to those with normal cognition reflects cell-specific heterogeneity in XCI or reflects a fundamental shift in XCI for these genes in AD, so additional work on XCI for these genes in the AD brain is warranted. In fact, our discovery of sex-biased expression pattern change for selected XCI genes during AD dementia is in line with a very recent study^92^ which revealed aging remodels transcription of the inactive X (Xi) and active X (Xa) across hippocampal cell types in female mice. The study observed that some genes on Xi underwent activation whereas some others showed new escape with aging, both in a cell-type-specific manner. It is safe to say that AD dementia-induced differential expression of X chromosome genes in certain brain cell types might contribute to sex differences of disease pathogenesis.

The most interesting finding of all sex chromosomal genes is *ASMTL-AS1* from CUX2-neurons, a long non-coding RNA (lncRNA) that resides in the PAR (i.e. pseudoautosomal region) of the X chromosome. Surprisingly, it showed male-biased expression among those with AD dementia (FDR<0.001) in CUX2-excitatory neurons and was not differentially expressed by sex among those with normal cognition (FDR=0.4). Upon further investigation, this male-biased expression seen in AD was caused by a severe reduction of *ASMTL-AS1* expression in excitatory neurons among females with AD (FDR=0.01). The XCI escape pattern of *ASMTL-AS1* is unknown, however its antisense gene *ASMTL* is a known XCI escapee^93^. Dysregulated expression of *ASMTL-AS1* was reported in several cancers^94–96^, but no evidence has yet linked it to neurodegenerative diseases. In fact, the role of lncRNA genes in non-tumorous diseases is just beginning to emerge, especially in AD^97,98^. A recent report discussed how a one copy deletion of a lncRNA gene, *CHASERR*, caused a neurodevelopmental disorder by increasing expression of an adjacent gene *CHD2*^99^.

### Damage-related stress-response pathways show notable sex-specific alterations in AD

We observed sex-specific enrichment for a number of metabolism-related and stress-response related pathways, including Hypoxia, DNA Repair, Oxidative Phosphorylation, and UV Response, all of which are related to well-known AD risk factors^100^. Stress-response gene signatures are upregulated in the AD brain across multiple cell types, especially in latest stages of pathology^50,100–102^, supporting our findings. Differences in both metabolism and stress-response have been highlighted as key contributors to sex differences in AD in a recent review due to the striking sex differences in the alterations of these key pathways over the course of aging and AD^103^.

Among these top enriched pathways for both sexes, Oxidative Phosphorylation (OXPHOS) is of particular interest. OXPHOS is another pathway that had the leading edge of enrichment with opposite AD effect genes in different sex. It was enriched in both endothelial cells and astrocytes for tau (protective genes enriched in females and risk genes enriched in males). OXPHOS is a process that takes place in mitochondria which produces ATP to meet cell’s energy demand through oxidizing NADH or FADH. Mitochondrial dysfunction and oxidative damage in AD patients have been documented as early as 1990s^104,105^. The mitochondrial cascade hypothesis for AD, which was first proposed in 2004^106^, posits that AD endophenotypes, such as Aβ production and tau phosphorylation, arise as a consequence of the compensation response to mitochondrial function and durability changes during aging^107^. Indeed, multiple studies showed AD pathology promoted dysfunctional mitochondria^108–111^. Our work suggests that sex differences may play a crucial role in this pathway.

One of the mechanisms by which abnormal OXPHOS (oxidative stress) contributes to AD is through damaging the blood-brain barrier (BBB), promoting the release of proinflammatory cytokines, and upregulating genes and proteins which can affect BBB permeability^112–114^. BBB dysfunction can expedite tau formation and dispersion^115^, and tau alone can initiate BBB breakdown with tau-induced glial activation and endothelial damage^116,117^. Endothelial cells and astrocytes are two essential components of BBB, which mediate the formation of tight junctions that critically affect the BBB function. Sex difference in BBB strength, shear stress response, vascular function, and metabolism has been observed, and multiple potential mechanisms including the influence from sex hormones have been proposed^118^. The interesting tau-related OXPHOS enrichment patten we observed in both endothelial cells and astrocytes might indicate BBB dysfunction through dysregulation of OXPHOS. In males, since OXPHOS enrichment was driven by genes that related to higher levels of tau pathology, the upregulation of OXPHOS in these cell types may trigger more tau buildup. In females, OXPHOS enrichment was driven by genes that were related to lower levels of tau pathology, so the upregulation of the same OXPHOS in females may serve as a protective mechanism against tau pathology.

### Sex-influenced and disease-vulnerable cell subpopulations

Recent studies^25,26^ from both ROS/MAP and SEA-AD cohorts demonstrated SST GABAergic neurons are the most vulnerable and the earliest affected neuronal subtypes in AD, and their downregulation in AD brains is evident, validated in both DLPFC and middle temporal gyrus (MTG) brain tissues. In our research, we observed that males had a significantly higher fraction of SST inhibitory neurons compared to females, and a nominally significant lower SST neuron fraction in AD dementia group compared to cognitive normal controls for both sexes, with a slightly larger disease-related reduction among females (-1.5%) compared to males (-0.6%). However, neither cell fraction nor weighted linear regression analysis of cell abundance find robust evidence of statistically significant sex interaction effect. That said, the SST neurons remain an interesting focus for future cell-specific work to clarify if there are sex differences in the loss of this important neuron population.

In addition, a recent study^26^ from SEA-AD demonstrated that certain oligodendrocyte subpopulations (“Oligo_2” and “Oligo_4”) were the most vulnerable cell types during AD pathogenesis and progression. Our cell fraction results showed that females had a higher fraction of enhanced-translation Oli.6 and Oli.12, while males had a higher fraction of Oli.3 and Oli.9. However, due to contradicting subtype classifications between studies, it is difficult to tell whether these sex differences map onto the previously reported disease alterations in oligodendrocytes. Again, our cell composition models did not show robust evidence of a statistically significant sex interaction effect with these oligodendrocyte subpopulations.

Most importantly, our study identified two more disease-associated, disease-vulnerable cell subpopulations in DLPFC, aSMC and Exc.3. Previous study^119^ observed a strong loss of brain vascular nuclei - across endothelial cells, SMCs, pericytes and fibroblast-like cells, though no association with AD diagnosis or endophenotype was detected in their dataset. A recent study^31^ investigating the relevance of fraction quantitative trait locus (fQTL) to AD risk using both snRNA-seq and bulk RNA-seq DLPFC data from ROS/MAP cohorts discovered one known AD risk SNP, rs5011436, that coincided with the fQTL of excitatory neuron 3 (Exc.3). Specifically, the major allele rs5011436^A^, the risk allele for AD, is negatively associated with Exc.3 fraction. This SNP resides within *TMEM106B*. The authors further demonstrated rs5011436 not only was an fQTL but also had a cis-eQTL effect on *TMEM106B* expression. Our discovery bolsters evidence that Exc.3 cell composition is altered in AD dementia. As for the aSMC subpopulations, more evidence is needed to understand its specific role in AD pathogenesis. aSMCs were showed to display profound organotypicity^120^, which suggested their functions being highly dependent on tissue environment. This might hint their modulation potential in disease.

### Limitations and future work

This manuscript included multiple strengths. The large sample size, longitudinal data prior to death from ROS/MAP, comprehensive clinical and neuropathological characterization of participants, independent replication, and deep sequencing provided the most comprehensive molecular profile of sex differences in the AD brain to date. Yet, the work is not without limitations. While we were able to replicate some effects, differences in sample size and cohort characteristics may have prevented additional replications. For example, basic demographic differences between the two cohorts, such as age at death, education, and sex distribution were notable. The SEA-AD cohort also included a greater proportion of advanced Braak stage patients (Braak 5 or 6) compared to the ROS/MAP cohorts. In terms of the sequencing data, they were prepared and generated in different settings and were processed based on different pipelines, which not only affected the discovery power but also created unique technical noise during replication. And finally, drastic differences in cell taxonomy and clustering algorithms used to produce cell types from ROS/MAP and SEA-AD cohorts prevented a rigorous assessment of cell identities and made cell-type based comparison very challenging. Hence, the low replication rate (0.4%) between these two studies was not surprising. More effort in the field should be focusing on data integration^26^ and phenotype harmonization^121^ to tackle abovementioned issues.

Another shortcoming of this study is that the ROS/MAP cohorts include more than twice as many females than males, providing more statistical power in analyses of females and likely contributing to the almost four-fold increase in female-specific gene associations compared to males. Our observations are also limited to non-Hispanic White individuals limiting our generalizability to other populations. Finally, our study only focused on DLPFC brain tissue. AD pathogenesis involves multiple brain regions and there are variations in AD pathology both spatially and temporally among different regions^122^. Our future effort should also include exploring other brain tissues as well as investigating evidence beyond transcriptomes, including protein translation, post-translational modifications, epigenetics, etc. to picture a broader overview of sex-specific effect of AD pathogenesis. Ultimately, experimental validation of some of these identified sex-specific gene effects will be crucial to move the field forward and realize the potential of sex-specific therapeutic interventions in AD.

## CONCLUSION

This manuscript has provided the most comprehensive analysis of sex-specific transcriptomic associations with AD endophenotypes at cellular resolution. Due to differences in sample size, we had more statistical power to identify true female-specific associations, and we were indeed able to identify and replicate multiple female-specific associations in neurons and astrocytes. Additionally, our results provide broad support for previous suggestions of enhanced neuroprotective associations in neurons for females, and enhanced risk associations in glial cells among males and females. While most of the female-specific associations we observed in our study were among neurons in relation to Aβ, most of the male-specific associations were observed among endothelial cells in relation to tau. Finally, our cell signaling analyses highlight multiple cell-cell interactions that appear to be altered in a sex-specific manner in AD. Together, these findings provide strong evidence of sex-specific transcriptomic drivers in the AD brain and provide a host of exciting sex-specific targets for intervention that can be validated and prioritized in future mechanistic studies.

## Data and code availability

Raw sequence and processed data of snRNA-seq from ROS/MAP study are available at Synapse (https://www.synapse.org/#!Synapse:syn31512863). FASTQ files of SEA-AD snRNA-seq are available through controlled access at Sage Bionetworks (accession no. syn26223298). Processed SEA-AD snRNA-seq are available through the Open Data Registry in an AWS bucket (sea-ad-single-cell-profiling) as AnnData objects (h5ad files) and viewable on CELLxGENE and Allen Brain Cell Atlas (https://portal.brain-map.org/atlases-and-data/bkp/abc-atlas). All codes used for analysis discussed in the manuscript are available upon request.

## Acknowledgement

We thank all participants and their families from each of the studies we mentioned for their generous donation and support for research. All sex difference analyses are supported through NIH funding R01AG061518. The results published here are in whole or in part based on data obtained from the AD Knowledge Portal (https://adknowledgeportal.org).

Study data of ROS/MAP were provided by the Rush Alzheimer’s Disease Center, Rush University Medical Center, Chicago. Data collection was supported through funding by NIA grants P30AG10161 (ROS), P30AG072975 (ROS), R01AG015819 (ROS/MAP; genomics and RNAseq), R01AG017917 (MAP), U01AG46152, U01AG61356, R01AG030146, RF1AG57473 (snRNA-seq), the Illinois Department of Public Health (ROS/MAP). Additional phenotypic data can be requested at www.radc.rush.edu. The Seattle Alzheimer’s Disease Brain Cell Atlas (SEA-AD) consortium is supported by the National Institutes on Aging (NIA) grant U19AG060909. Study data were generated under the support of in part by the UW (University of Washington) Alzheimer’s Disease Research Center (NIA P30AG066509), the Adult Changes in Thought (ACT) Study (NIA U19AG066567), and the Nancy and Buster Alvord Endowment (to C.D.K.). Data collection for this work was additionally supported, in part, by prior funding from NIA (P50AG005136; U01AG006781).

## Author Contributions

Y.W. and T.J.H. designed the study and wrote the manuscript. Y.W. performed all the analyses. K.T., M.G., D.K, P.L.D.J, V.M, J.A.S, D.A.B. generated data, aided in the interpretation of results, and provided critical revision of the manuscript. A.D., C.D.K., L.D. contributed to study design and provided critical revision of the manuscript. All authors approved the final manuscript.

## Conflict of Interest

T.J.H. serves on the Scientific Advisory Board for Vivid Genomics, serves as Deputy Editor for Alzheimer’s & Dementia: Translational Research and Clinical Intervention, and as Senior Associate Editor for Alzheimer’s & Dementia. All other authors have no competing interests.

## Extended data figures and tables

**Extended Data Fig 1:**
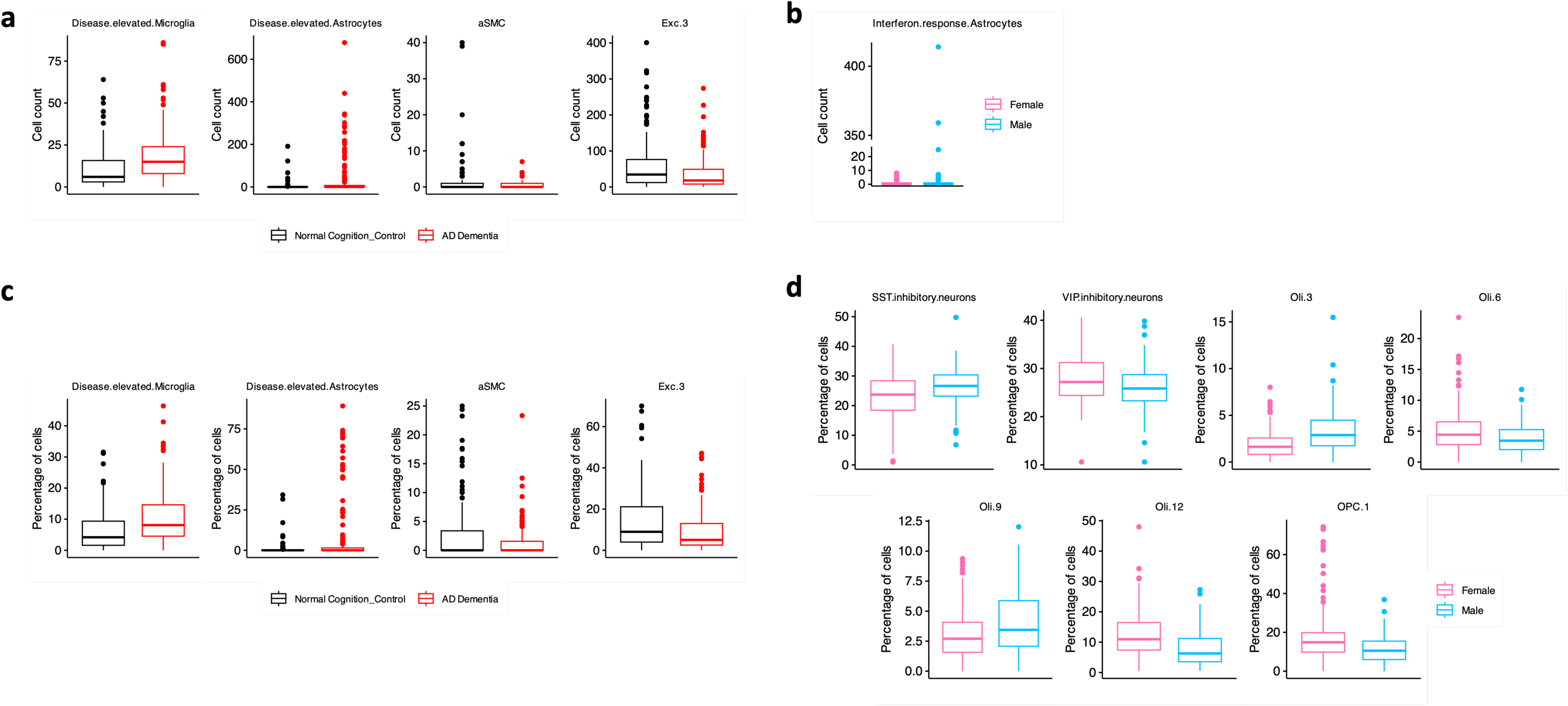
ROS/MAP Cell Composition influenced by sex or AD dementia diagnosis. a, Boxplots showing the four cell subtypes that are influenced by AD dementia diagnosis. b, Boxplots showing the cell count for interferon-response astrocytes that are influenced by sex. c, Boxplots showing fraction (percentage) of the four cell subtypes that are influenced by AD dementia diagnosis. d, Boxplots showing fraction (percentage) of the seven cell subtypes that are influenced by sex.

**Extended Data Fig 2:**
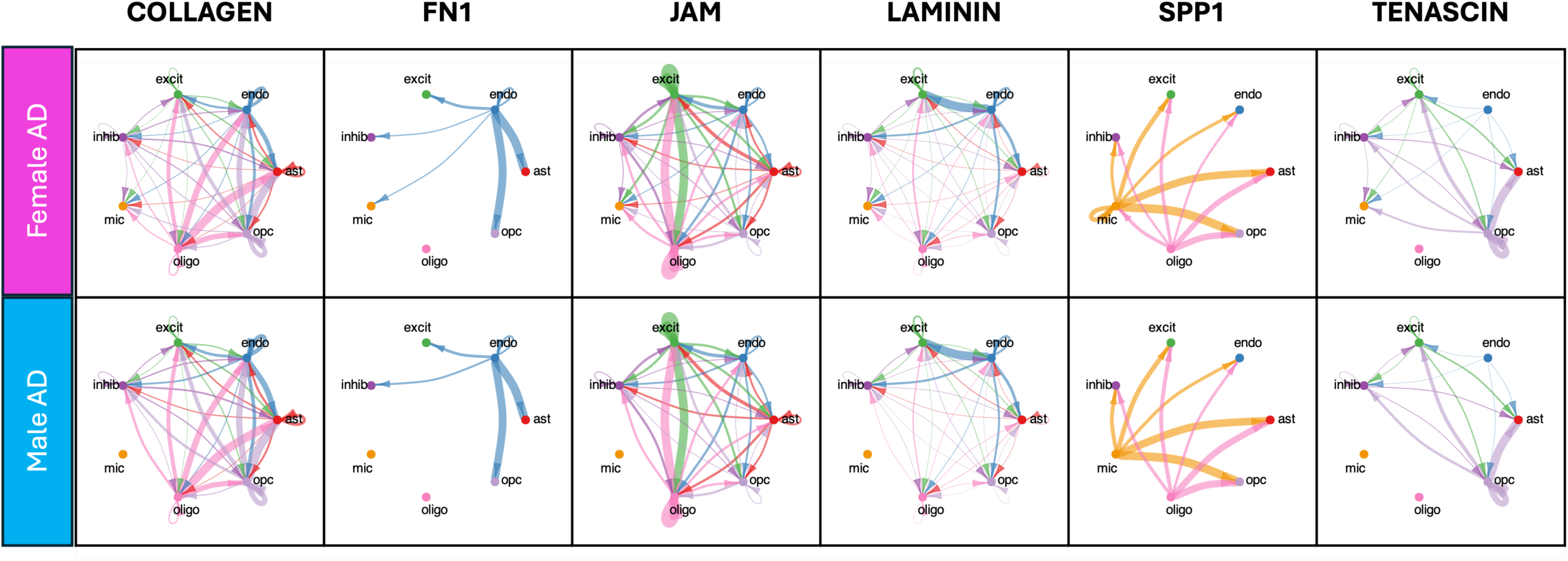
Six microglial-incoming signals comparison. Chord diagram showing six signaling pathways in females and males with AD dementia. Males with AD dementia lacked meaningful incoming signals of each of these signals to microglia compared to females with AD dementia. Cell type abbreviation: ast, astrocytes; excit, excitatory neurons; inhib, inhibitory neurons; mic, microglia; oligo, oligodendrocytes; opc, oligodendrocyte precursor cells; endo, endothelial cells. Each information flow represents combined interactions from all enriched ligand-receptor pairs.

## Notes

https://www.synapse.org/#!Synapse:syn31512863

https://portal.brain-map.org/atlases-and-data/bkp/abc-atlas

### Competing Interest Statement

T.J.H. serves on the Scientific Advisory Board for Vivid Genomics, serves as Deputy Editor for Alzheimers & Dementia: Translational Research and Clinical Intervention, and as Senior Associate Editor for Alzheimers & Dementia. All other authors have no competing interests.

